# Merits and challenges of plasma proteomics on association replicability

**DOI:** 10.1101/2025.09.01.673490

**Authors:** Zeyu Jiao, Yixin Zhang, Yinglei Lai, Jujiao Kang, Liang Ma, Wei Zhao, Jia You, Wei Cheng, Jianfeng Feng

## Abstract

Developments in proteomic platforms have enabled the generation of large-scale high-throughput plasma proteomics data [1–3]. With recent breakthroughs in AI modelling, these data have significantly enhanced our understanding of molecular mechanisms underlying human behaviors and diseases [4–6]. However, the replicability of associations between plasma proteomics and phenotypes remains underexplored. Here, we systematically assessed the replicability of associations with recent plasma proteomics data in the UK biobank. Over 75% of cognitive function and mental health traits demonstrated high overall (proteomics-wide) replicability when brain-related traits were considered as phenotypes. Although mean cortical thickness (CT) as phenotype exhibited clearly reduced replicability, total cortical surface area (CSA) and cortical volume (CV) showed high overall replicability across hemispheres and over twenty brain regions. In comparative multi-omics analyses based on the same cohort of participants, proteomics outperformed genomics across all brain-related traits, and exceeded metabolomics for over half of traits where metabolomics also exhibited high overall replicability. Furthermore, we developed a predictive framework to estimate the replicability for potential future proteomics panels based on the crucial influential factors including dilution level, proportion of samples below the limit of detection (LOD), and sample size. Moreover, we constructed an individual replicability index for proteins and identified eleven proteins with highly replicable associations across cognitive function and mental health traits, which was in line with the recent identifications of pleiotropic proteins in large-scale population studies. Collectively, our results revealed the challenges in the association replicability of plasma proteomics under reduced data quality (from “Explore” to “Expansion” assay panels), and we further explored how to sustain high replicability in potential future panels. Fundamentally, our findings affirm the merits of plasma proteomics: this molecular omics platform enables highly replicable associations for mapping biomedical signatures.

## Introduction

The development of high-throughput proteomics platforms and recent breakthroughs in artificial intelligence (AI) have enhanced our understanding of the molecular mechanisms underlying various human behaviors and diseases [1–6]. In recent years, association analyses remain fundamental in large-scale high-throughput studies, yet replicability challenges arise in these studies [7, 8]. For example, the replicability issues of genome-wide association studies (GWAS) have garnered a considerable attention [9]. Furthermore, the replicability of brain-wide association studies (BWAS) was also assessed based on large-scale neuroimaging data in recent studies [8, 10]. Given the continuous expansion for proteomics-based association studies (PBAS), there is still a lack of systematic replicability investigations. Moreover, the prospect of diagnosing complex conditions through accessible blood tests represents a promising change in health care, which offers an early-stage alternative to costly and invasive procedures [11]. Plasma proteomics has been demonstrated as a cutting-edge approach for the development of blood tests [12], although it has not yet been systematically compared to other blood-based platforms (e.g., genomics and metabolomics) in terms of association replicability.

Several criteria have been developed to quantify the replicability of associations, generally based on statistical significance [13, 14], rank correlation [15], or directional consistency (DC) [16, 17], etc. Among these, the DC criterion is specifically developed based on the consideration of both high-throughput settings and consistencies of association directions (i.e., overall replicability), thereby useful in the replicability assessment for association analysis findings [16–19]. Here, we adopted the DC criterion for our systematic assessment of replicability in PBAS. Notably, the overall replicability (based on directional consistency) may be impacted by influential factors, such as the fraction of missing data, dilution level, proportion of samples below the limit of detection (LOD) and sample size [1, 7, 8, 20]. However, the impact of these factors on the overall replicability of PBAS has not yet been systematically characterized. Moreover, the recent plasma proteomics has been rapidly developed in both sample sizes and throughput volumes [21]. A comprehensive understanding of these influential factors is essential for estimating the association replicability of potential future proteomic panels. Here, we intend to address these issues in this study.

Recently, the identifications of pleiotropic proteins have been gaining a significant attention to understand systems biology and complex diseases [22]. These proteins can substantially lower the high expenses of broad proteomic screening and facilitate their translation into potential clinical targets [6]. Notably, an individual-level assessment of association replicability for each protein provides an essential contribution to the development of pleiotropic proteins [15, 16]. Therefore, in this study, we also aim to develop a procedure for this purpose.

To our knowledge, we were among the first to conduct a comprehensive investigation of the DC-based replicability for associations in plasma proteomics. Based on this investigation at the individual level for proteins, our study provided an alternative perspective to the discovery of pleiotropic proteins, which could facilitate the development of practically useful biomarkers. Based on the investigation of overall replicability, we enhanced our confidence in the current development and applications of high-throughput plasma proteomics, and highlighted the challenges along with the growth of throughput volume and scale. As the development of future proteomic platforms would be impacted by certain crucial factors, our study provided a related insight, and also addressed the prediction of association replicability for potential future plasma proteomic platforms. Fundamentally, our study contributed to a systematic understanding of the merits of plasma proteomics on association replicability. (Details of study overview please see Figure 1.)

**Figure 1.**
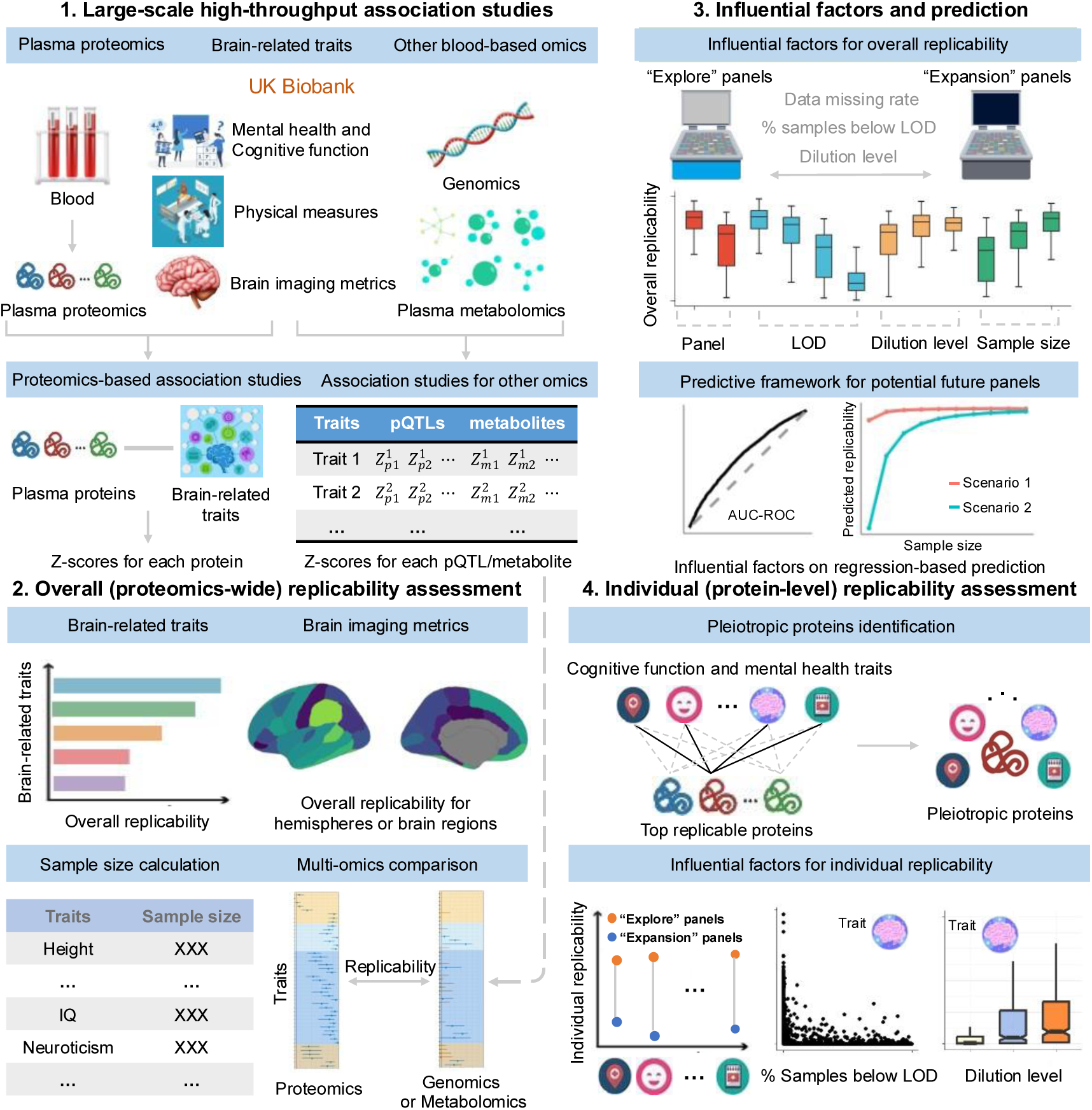
| Overview of the study design and analyses. Part 1. Large-scale association studies utilizing UK Biobank plasma proteomics, phenotypes, genomics, and plasma metabolomics. The full list of brain-related traits within physical measures, cognitive function, mental health and brain region names for imaging metrics could be found in Supplementary Table 1. **Part 2.** Overall (proteomics-wide) replicability assessment for PBAS, sample size calculation and multi-omics comparison. **Part 3**. Characterization of influential factors on overall replicability and development of a predictive framework for potential future panels. **Part 4.** Individual (protein-level) replicability assessment with pleiotropic proteins identification and influential factors characterization.

## Results

### Overall replicability for PBAS

The simulation results, summarized in Supplementary Materials and Figure S1, indicated that the MMRA approach could accurately assess the replicability of PBAS results.

As shown in Figure 2a and 2b, for all physical measures including body mass index (BMI), height, weight, diastolic blood pressure (DBP), systolic blood pressure (SBP), speech-reception-threshold (SRT) estimate and pulse wave arterial stiffness index (ASI), a median overall irreplicability quantity 𝜌_*IR*_ < 0.05 was consistently observed for each trait. For all 31 traits within cognitive function and mental health, the median overall irreplicability quantity 𝜌_*IR*_ < 0.05 was observed for 24 traits (77.4%) indicating high levels of overall replicability. For cognitive function, the median 𝜌_*IR*_ ≥ 0.05 was observed in two traits. The median 𝜌_*IR*_ was 0.429 for the symbol digit substitution test and 0.539 for the paired associate learning test, with corresponding lower- and upper-quartiles (Q1-Q3) of 0.355-0.524 and 0.018-0.703, respectively. For the 24 traits under mental health, only 5 traits exhibited low overall replicability levels, with a median 𝜌_*IR*_ ≥ 0.05. In these five traits, such as the work satisfaction and family relationship satisfaction, the values of median 𝜌_*IR*_ were relatively higher (The median 𝜌_*IR*_ was 0.505 for the work satisfaction and 0.568 for family relationship satisfaction). Please see Supplementary Table 2 for the details of assessed median 𝜌_*IR*_ with corresponding lower- and upper-quartiles (Q1-Q3) for traits within physical measures, cognitive function and mental health. We further evaluated the influence of blood collection season and fasting time on PBAS replicability. Overall, these additional covariates had extremely limited influence on replicability, as shown in Figures S2 and S3.

**Figure 2.**
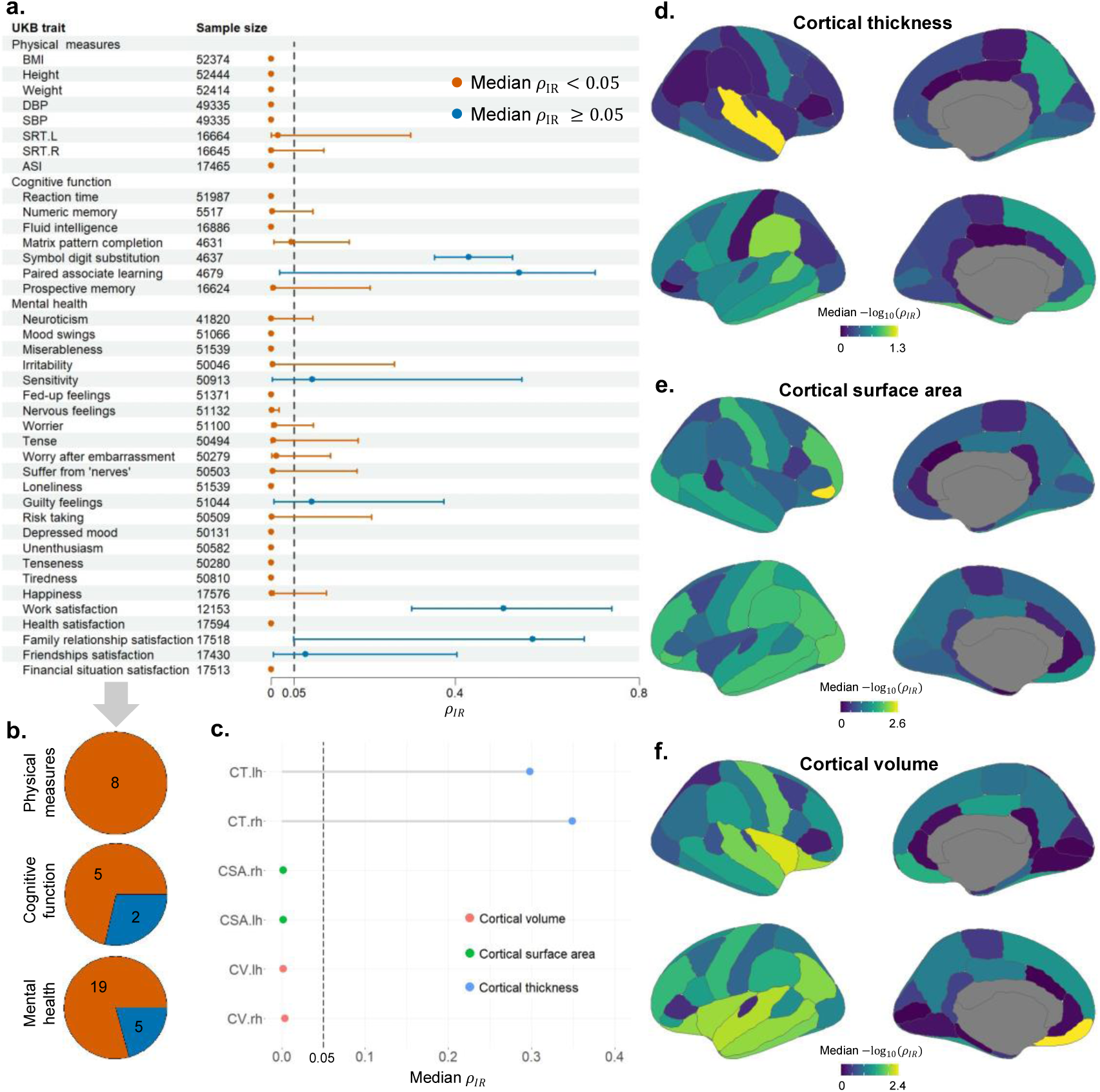
| Overall replicability assessment for brain-related traits. **a,** Median overall irreplicability quantity (𝜌_*IR*_) with lower- and upper-quartiles (Q1-Q3) from 1,000 random subsampling times for traits in physical measures, cognitive function, and mental health, alongside corresponding sample sizes. **b,** Number of traits within physical measures, cognitive function, and mental health with median 𝜌_*IR*_ < 0.05 versus 𝜌_*IR*_ ≥ 0.05. **c,** Median 𝜌_*IR*_ for mean cortical thickness (CT), total cortical surface area (CSA), and cortical volume (CV) in both hemispheres. **d–f,** Regional median − log_10_(𝜌_*IR*_) for (**d**) mean CT, (**e**) total CSV, and (**f**) total CV.

For brain imaging metrics, as shown in Figure 2c, the median 𝜌_*IR*_ values were 0.2983 and 0.3492 with the Q1-Q3: 0.0529-0.5326 and 0.0776-0.5346 for mean CT in left and right hemispheres, respectively. However, total CSA and CV showed clearly higher overall replicability than mean CT. The median overall irreplicability quantity 𝜌_*IR*_ values for total CSA in left and right hemispheres reached 0.0017 and 0.0014 while the related Q1-Q3 was 0.0001-0.0743 and 0.0001-0.1071, respectively. For total CV in left and right hemispheres, the median 𝜌_*IR*_ values were 0.0017 and 0.0036 while the related Q1-Q3 was 0.0002-0.1298 and 0.0002-0.1323, respectively. These results indicated a high level of overall replicability for brain imaging metrics. Please see Supplementary Table 3 for the details of assessed median 𝜋_*R*_, 𝜋_*IR*_or 𝜌_*IR*_ for total CSA, total CV and mean CT in left and right hemispheres.

No regions were identified for region-wide mean CT with median 𝜌_*IR*_ < 0.05 (Figure 2d). For region-wide CSA versus plasma proteins, when using the threshold of median 𝜌_*IR*_ < 0.05, twenty-one brain regions were identified in our results (Figure 2e). The top three regions were the right pars orbitalis gyrus (median 𝜌_*IR*_ = 0.0024), followed by the left lateral occipital gyrus (median 𝜌_*IR*_ = 0.0106) and the right rostral middle frontal gyrus (median 𝜌_*IR*_ = 0.0113). For CV, twenty-three regions were identified with median 𝜌_*IR*_ < 0.05 (Figure 2f). The top three regions were the left medial orbitofrontal gyrus (median 𝜌_*IR*_ = 0.0037), followed by the right insula gyrus (median 𝜌_*IR*_ = 0.0050) and the left superior temporal gyrus (median 𝜌_*IR*_ = 0.0069). (Please see Supplementary Table 4 for the details of median 𝜌_*IR*_ for region-wide CSA, CV and CT.) The potential explanations for the observed differences in overall replicability levels between total CSA/CV and mean CT were given in supplementary materials and Figure S4.

### Multi-omics overall replicability comparison

As shown in Figure 3a and 3b, compared to pQTLs, association analysis results based on proteomics data exhibited notably lower 𝜌_*IR*_ values, indicating higher overall replicability. This comparison was based on the same cohort of participants. For all 31 traits within the categories of cognitive function and mental health, we could also observe the median overall irreplicability quantity 𝜌_*IR*_ < 0.05 on 23 traits (74.2%), indicating high levels of overall replicability for proteomics data (Figure 3b). For genetics-based association analyses, a high replicability level (median 𝜌_*IR*_ = 0.00082) was observed only for height. For brain-related measures, relatively low overall replicability levels (median 𝜌_*IR*_ ≥ 0.05) were observed in genetic-based analyses.

**Figure 3.**
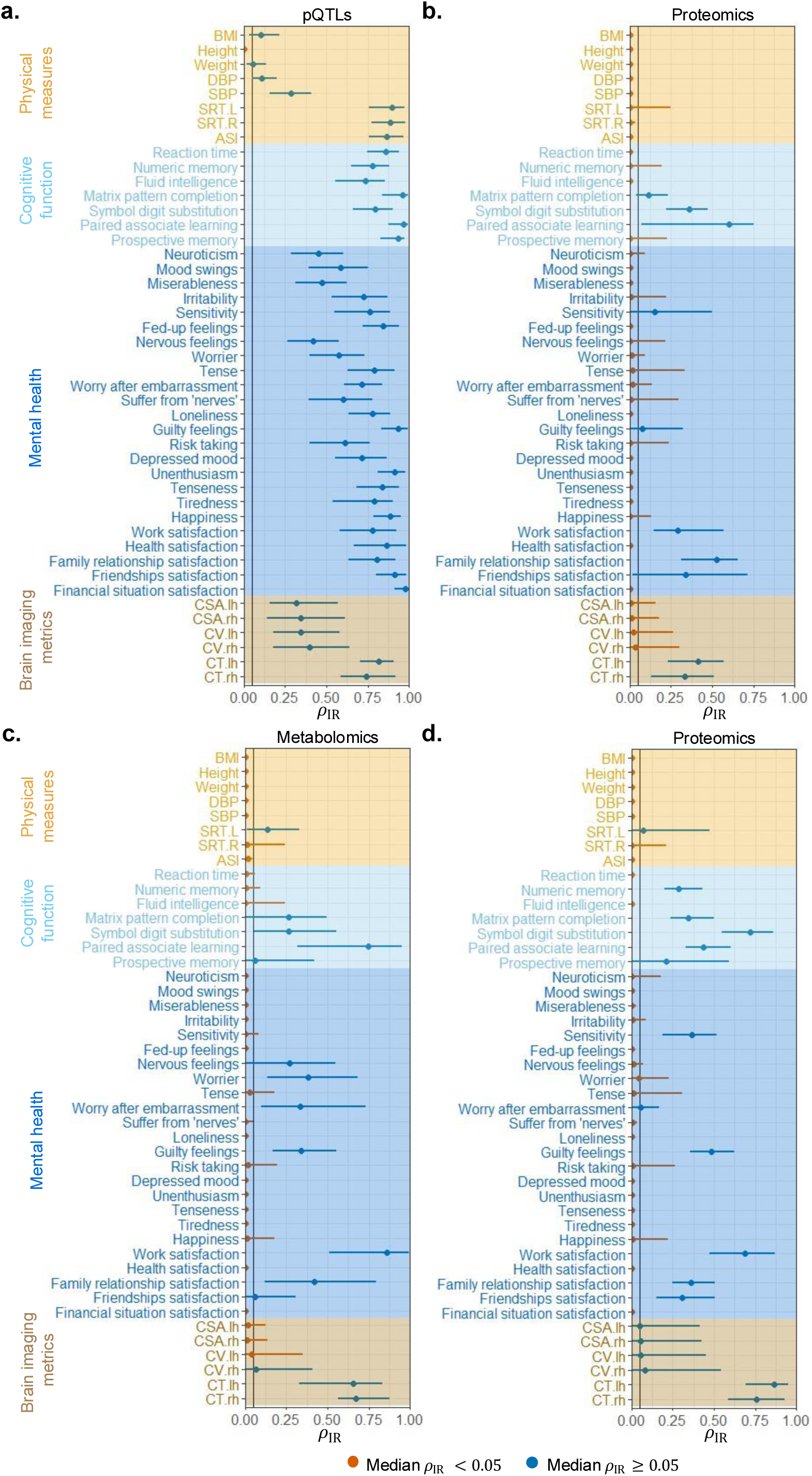
| Comparative overall replicability across multi-omics. Median overall irreplicability quantity (𝜌_*IR*_) and the lower- and upper-quartiles (Q1-Q3) from 1,000 random subsampling times are shown. **a, b,** Comparison between genetics-based associations from pQTLs and PBAS. **c**, **d**, Comparison between metabolomics-based associations and PBAS. Traits are grouped into physical measures, cognitive function, mental health, and brain imaging metrics.

Based on the same cohort of participants, the overall replicability comparison of metabolomics-based association studies versus PBAS was presented in Figure 3c and 3d. Here, we observed the median overall irreplicability quantity 𝜌_*IR*_ < 0.05 in 7 out of 8 physical measures. Moreover, among the 31 traits related to cognitive function and mental health, 20 traits (64.5%) exhibited the median 𝜌_*IR*_ < 0.05. For brain structure imaging metrics, the median 𝜌_*IR*_ < 0.05 was observed for total CSA in both hemispheres and total CV in the left hemisphere. While these results indicated that metabolomics-based association studies also demonstrate high levels of overall replicability, PBAS exhibited even higher overall replicability for all 8 physical measures and over half of brain-related phenotypes (i.e., 17 out of 31 traits related to cognitive function and mental health shown in Supplementary Table 5). Further details of overall replicability comparisons of genetics versus proteomics and metabolomics versus proteomics were also displayed in Supplementary Table 5.

### Influential factors for overall replicability

The proteomics data quality difference between Panels 1 (“Explore 1,536 assay panels”) and 2 (“Expansion 1460 assay panels”) may lead to different overall replicability for PBAS. Panel 2 showed a clearly higher missing rate: the median missing rate was 0.169 while Q1-Q3 was 0.152-0.175. Panel 1 displayed impressively low missing rate: the median missing rate was 0.0274 while the related Q1-Q3 was 0.0231-0.0423. Moreover, the category with least-abundant (1:1) dilution section and more than 50% of samples below LOD in Panel 2 contained clearly more proteins than that in Panel 1 (see Supplementary Table 6 for details). When phenotypes fluid intelligence (Figure 4a) and neuroticism (Figure 4b) were considered, we observed clearly lower overall replicability for Panel 2.

**Figure 4.**
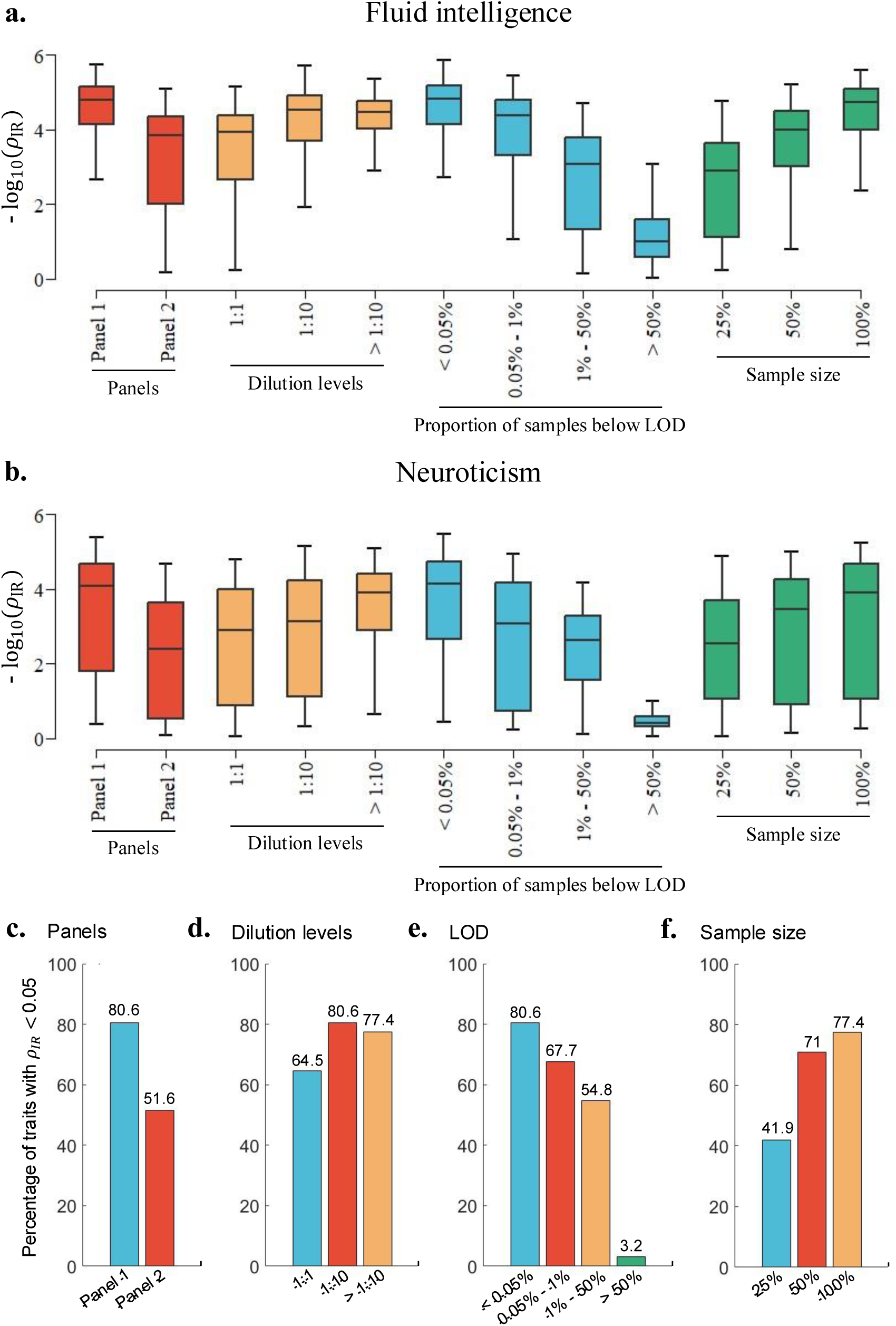
| Characterization of influential factors for overall replicability. **a, b,** The overall replicability (𝜌_*IR*_) for (**a**) fluid intelligence and (**b**) neuroticism with different panels, dilution levels, the proportion of samples below LOD and study sample size. Each box plot represents the – log_10_ (𝜌_*IR*_) from 1,000 random subsampling times. The center line in each box represents the median; the lower and upper hinges represent the 25th and 75th percentiles, respectively; the whiskers represent 1.5× the lower- and upper-quartiles. **c-f**, Percentage of traits with median 𝜌_*IR*_ < 0.05, summarized separately for (**c**) panels, (**d**) dilution levels, (**e**) the proportion of sample below the LOD, and (**f**) sample sizes.

We then investigated the impact of proteomics data quality for overall replicability and the results were also demonstrated in Figure 4. Considering fluid intelligence and neuroticism as examples, we observed that the least-abundant (1:1) dilution section exhibited relatively higher overall irreplicability quantity 𝜌_*IR*_ values compared to the moderate-abundant (1:10) and more-abundant (1:100 to 1:100000) dilution section. Across four categories stratified by the proportion of samples below the LOD (<0.05%, 0.05%-1%, 1%-50%, >50%), overall replicability declined as the proportion of samples below the LOD increased. A downward trend was observed for overall irreplicability quantity 𝜌_*IR*_ with increasing sample size.

For additional traits, a clear decline in overall replicability was observed in Panel 2, lower dilution levels, higher proportions of samples below the LOD and smaller sample size (Figure S5-S8). Based on Panel 2, only 51.6% of traits related to cognitive function and mental health showed high overall replicability (median 𝜌_*IR*_ < 0.05), compared to 80.6% based on Panel 1 (Figure 4c). Based on the least-abundant dilution section (1:1), 64.5% of cognitive and mental health-related traits exhibited high overall replicability, compared to 80.6% and 77.4% of traits based on the moderate-abundant (1:10) and more-abundant (1:100 to 1:100,000) dilution sections, respectively (Figure 4d). Based on the categories of proteins with more than 50% of samples below the LOD, only 3.2% of traits demonstrated high overall replicability; this percentage was substantially lower than those based on the other three LOD-based categories (Figure 4e). When the sample size was reduced to approximately 50% and 25% of the original dataset, 71% and 41.9% of traits respectively exhibited high overall replicability, compared to 77.4% based on the original dataset (Figure 4f). Detailed results were provided in Supplementary Table 7.

### Replicability or potential future panels

When the fluid intelligence and neuroticism were considered as phenotypes in our logistic regression model, larger sample size and lower proportion of samples below the LOD showed positive contributions while negative for lower abundant of dilution level (Figure S9); and AUC-ROC of 0.62 and 0.58 were achieved, respectively (Figure S10). According to the difference between Panel 2 versus Panel 1, we would consider a potential future panel with lower dilution level and higher proportion of samples below the LOD. For the hypothetical examples presented in Figure S10, increasing sample size would generally improve the predicted probability of overall replicability, though the improvements may be modest for certain phenotypes such as neuroticism.

### Pleotropic proteins identification

The top ten proteins with the highest individual replicability were identified as being highly replicable in their associations with each trait. The full list of top ten proteins for each PBAS scenario could be found in Supplementary Table 8. GDF15 was identified as a pleotropic protein (See Methods section for details) in both mental health (in 14 traits) and cognitive function (in 3 traits). The proteins ASGR1, PIGR, PLAUR, and PRSS8 were identified as pleotropic proteins in mental health, appearing in 10 to 14 traits; and the proteins APOF, CCL20, CDCP1, GGH, MZB1 and TFF1 were identified as pleotropic proteins in cognitive function (Figure 5a).

**Figure 5.**
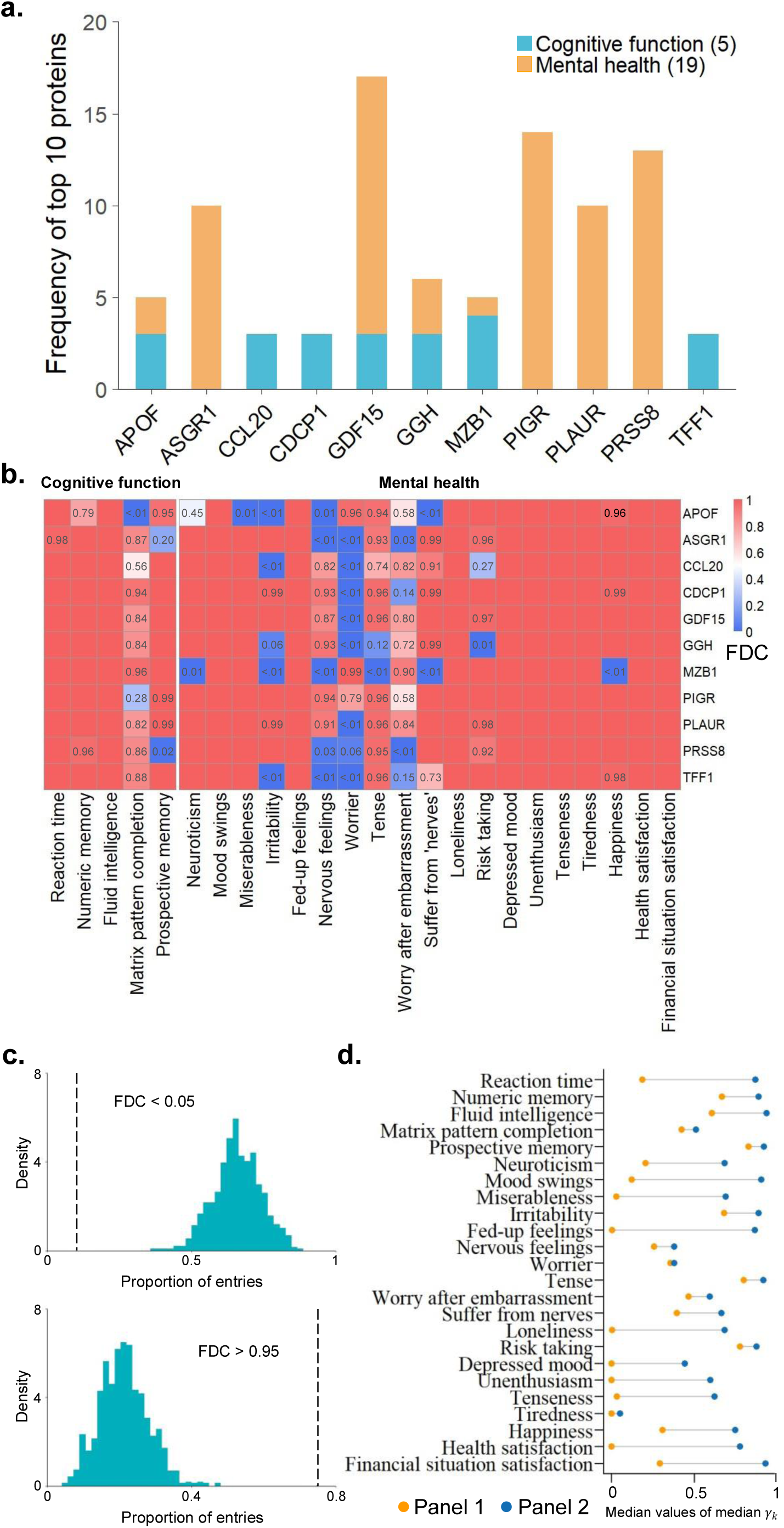
| Individual-level replicability assessment. **a,** Pleiotropic proteins and their observed frequency of being ranked among the top 10 across traits in cognitive function and mental health. **b,** FDC for each pleotropic protein across traits. Blank cells indicated FDC > 0.99. **c,** Proportion of entries with extreme FDC values (upper: FDC < 0.05, lower: FDC > 0.95) for pleiotropic proteins versus randomly selected proteins. Dashed lines indicated the observed proportions of pleiotropic proteins. **d,** Median values of median individual irreplicability quantity (𝛾_𝑘_) between Panel 1 and Panel 2 for each trait within cognitive function and mental health.

As shown in Figure 5b, an FDC > 0.99 (fraction of directional consistency, see Methods section for details) was observed in all 1,000 random subsampling for at least one trait in the cognitive function and mental health for each pleotropic protein. Many traits exhibited FDC > 0.95 for all these pleotropic proteins (such as reaction time, fluid intelligence in the cognitive function, as well as mood swings, fed-up feelings, loneliness, depressed mood, unenthusiasm, tenseness, tiredness, health satisfaction and financial situation satisfaction in the mental health). For certain traits, such as worry and worry after embarrassment, many pleiotropic proteins showed FDC < 0.05. Nevertheless, the protein MZB1 demonstrated high FDC values of 0.99 and 0.90 for worry and worry after embarrassment, respectively. In summary, for every trait considered, there was at least one pleotropic protein exhibited an FDC > 0.9.

To support the significance of eleven pleotropic proteins for cognitive function and mental health, we randomly selected eleven proteins from the remaining 2,909 proteins and calculated the FDCs. For each set of randomly selected eleven proteins, we calculated the proportion of entries (total 11 × 24 proteins·traits entries) with FDC < 0.05, indicating relatively high individual irreplicability. This selected procedure was randomly repeated 1,000 times. Similar results were also calculated for the proportion of entries with FDC > 0.95. The corresponding distributions were shown in Figure 5c. Compared to the random selections, the set of eleven pleotropic proteins exhibited significantly fewer proportion of entries with FDC < 0.05 and substantially more proportion of entries with FDC > 0.95.

### Influential factors for individual replicability

The individual replicability level for Panel 2 was clearly lower than Panel 1 (Figure 5d). For example, for mood-related phenotypes such as depressed mood and unenthusiasm, we observed low median 𝛾_𝑘_ values of 0.001 and 5.24×10^-4^ for all the proteins in Panel 1; however, these values were 0.443 and 0.596 for Panel 2. Please see Supplementary Table 9 for the details of median 𝛾_𝑘_ for Panels 1 and 2. In addition, the individual irreplicability quantity 𝛾_𝑘_ of PBAS was also assessed for sub-panels in Panels 1 and 2, including Cardiometabolic, Inflammation, Neurology and Oncology. The assessment details can be found in the Supplementary Materials. For these four sub-panels, higher levels of individual replicability and a lower data missing rate were also observed in Panel 1 (Figure S11 and Supplementary Table 10).

When analyzing the impact of different dilution levels for individual replicability performance, we observed that the least-abundant dilution section (1:1) exhibited relatively higher median individual irreplicability quantity 𝛾_𝑘_ values for all traits compared to the moderate-abundant (1:10) and more-abundant (1:100 to 1:100000) dilution section (Figure S12). Detailed individual replicability assessments for each dilution section were provided in Supplementary Table 11. Additionally, we investigated the impact of the proportion of samples below the LOD on individual replicability. For each trait related to cognitive function and mental health, we found a significant negative relationship between individual replicability (median − log_10_ (𝛾_𝑘_)) and the proportion of samples with measurements below the LOD (Spearman’s 𝜌 < −0.1399, *P* < 10^−8^; Figure S13).

## Discussion

With the development of high-throughput proteomics platforms and AI modeling, the replicability of associations serves as a foundation for both biological discoveries and clinical translations [1–6]. While the advantages of plasma proteomics have been widely acknowledged [1–3], its association replicability remains a critical yet underexplored area. In order to further understand the merits and challenges in plasma proteomics, we provided a comprehensive investigation of the DC-based replicability for associations. Our work highlighted three key insights: (1) We demonstrated the high overall association replicability of plasma proteomics, which underlined its advantages versus genomics and metabolomics platforms; (2) We assessed crucial influential factors for association replicability, and developed a predictive framework according to potential future challenges along with the growing throughput of proteomics; and (3) Based on an individual-level replicability index and its related evaluation procedure, we identified eleven replicable pleiotropic proteins for cognitive function and mental health.

A key strength of our study lies in its comprehensive evaluations, which provide the depth and breadth in assessing the association replicability of proteomics. Across a diverse range of phenotypes, including physical measures and brain-related traits (cognitive function and mental health traits), we demonstrated the broad utility of plasma proteomics. For brain imaging measures, high overall replicability could also be observed for total CSA and CV in both hemispheres and over twenty brain regions. For mean CT in the proteomics-based association study, the proportion of no signals (i.e., true null hypotheses) was relatively higher than that for total CSA and CV (Figure S4). As the DC-based replicability assessment was influenced by the proportion of positive/negative signals and no signals, this could explain the lower overall replicability observed for mean CT. Likewise, a recent genomic study highlighted a comparatively larger genetic architecture for total CSA than that for mean CT [23]. Moreover, multi-omics data have emerged as a promising foundation for the development of blood tests which could enhance disease screening rates and facilitate early diagnosis [11, 24]. For example, molecular phenotyping based on genomic data facilitates early prediction and more accurate characterization of disease progression [25]. Metabolomics has emerged as a powerful approach for the identification of pre-disease states [26]. Furthermore, the prediction modeling based on plasma proteomics enables reliable estimation of 7-year to 10-year incidence risk for various common and rare diseases [27, 28]. By demonstrating the advantages of proteomics versus genomics and metabolomics in association replicability within the same cohort of participants, our study strengthened the confidence in both ongoing large-scale proteomic association studies and the desirable development of reliable blood-based diagnostic tools. Notably, to our best of knowledge, a large sample size and multi-phenotype dataset that contained matched plasma proteomics and transcriptomics was not available currently. The comparison between plasma proteomic and transcriptomics based on the same cohort of participants was not conducted.

While our findings affirmed the merits of plasma proteomics, we also faced challenges in replicability with the growth of throughput volume. To address this issue, we assessed the factors influencing the replicability of association findings (including missing rate, LOD, dilution level and sample size), and also developed a predictive framework for estimating the replicability of potential future assay panels. A crucial observation from our work is the concerning trend of declining data quality (proteins with higher missing rate, lower abundant dilution level and higher proportion of samples below LOD) in ‘Expansion 1460 assay panels’ versus ‘Explore 1,536 assay panels’. In high-throughput association studies, higher rates of missing data were shown to reduce the replicability of findings [7]. Furthermore, a recent study demonstrated the decrease of number of identified pQTLs versus the increase of dilution level or proportion of samples below the LOD [1]. Another study also shown the influence of these two factors on the correlations between the proteomics data generated from two different platforms [20]. In line with these previous studies, our results revealed the impact of these three factors (i.e., missing rate, LOD and dilution level) on the association replicability, and showed that the Expansion assay panels exhibited a relatively lower replicability than the Explore assay panels. Moreover, in our results, sample size was also identified as a crucial influential factor for replicability. Based on our predictive framework, we further demonstrated that increasing sample size can be a practically feasible way to sustain replicability for potential future proteomics panels. Adequate sample sizes were essential to ensure replicable results in high-throughput omics association studies [8, 12]. For instance, the results of GWAS with relatively small sample sizes generally failed to be replicated [29], while millions of samples were often required [30]. Similarly, growing concerns were raised on the sample size requirement for brain-wide association studies (BWAS), where thousands of samples are suggested to achieve satisfactory replicabilities [8]. In our recent study, a desirable overall replicability was achieved for physical measures when the sample size reached several hundred to a few thousand [17]. In this study, for plasma proteomics data, our results demonstrated that thousands of samples were sufficient to achieve a high overall association replicability for physical measures. For cognitive function, mental health, and brain imaging measures, a high overall association replicability was also achieved in most traits (21 out of 37 with sample size <10,000) when the sample size reached several thousands (Supplementary Table 12).

Our investigation included an individual level replicability assessment, which could also identify proteins with highly replicable associations across a scope of phenotypes. Eleven pleiotropic proteins were identified for cognitive function and mental health, including GDF15, ASGR1, PIGR, PLAUR, PRSS8, APOF, CCL20, CDCP1, GGH, MZB1 and TFF1. Among them, plasma GDF15 [31–35], PLAUR [36–38], CDCP1 [22, 39] and TFF1 [22] were identified as potential response biomarkers associated with cognitive function and mental health, and these proteins were further highlighted on the importance of pleiotropy in complex traits [22]. Moreover, ASGR1, PIGR, PRSS8 [36] and APOF [40] were identified as important protein biomarkers for mental diseases including depression, neurodegeneration and schizophrenia. Furthermore, CCL20 [41], GGH [42] and MZB1 [43] were associated with Alzheimer’s disease and reduced cognitive functions during the process of ageing. As pointed out by Topol [6], identifying a relatively small set of proteins with replicable associations could facilitate the development of targeted, low-cost assay panels for the proteomics-driven clinical translations.

Our study had several limitations. First, our analysis was primarily focused on a range of brain-related phenotypes. Although these traits are diverse and complex, the replicability levels and influential factors may not be directly generalized to other disease domains. Further investigations are warranted to a broader spectrum of human diseases. Second, our investigation was conducted based on the UK Biobank data. Consequently, our findings may not be representative to other populations. To address this limitation and the generalization of proteomic discoveries, it is necessary to conduct further investigations for diverse cohorts. Third, our results were based on the Olink high-throughput proteomics platform. Nevertheless, similar results may still be observed from proteomics data generated by other technologies.

Despite these limitations, our study provides evidences for the advantages of plasma proteomics in large-scale association studies. In summary, this study was among the first to provide a comprehensive DC-based assessment of the association replicability. Our study included the assessment at the overall and individual levels for association replicability. Our work further revealed the challenges for the future developments, and our analyses on influential factors and association replicability prediction provided a valuable contribution. Fundamentally, our findings affirmed that plasma proteomics was replicable in association analyses.

## Methods

### Study participants

This study included data from the UKB and integrated multiple data sources, including blood collection, imaging data, and various self-reported questionnaires [44, 45]. All participants provided explicit, written informed consent to the UKB. The UKB cohort received approval from the NHS National Research Ethics Service North West (reference number: 16/NW/0274).

Blood samples for proteomic analysis were collected and processed at Olink Analytical Services using the antibody-based Olink Explore™ Proximity Extension Assay. A total of 2,923 distinct proteins were measured, with stringent quality control procedures applied as outlined in the previous studies [1, 3]. Additional details regarding sample selection, processing, and quality control procedures were available in previous publications [1, 3]. The reported normalized protein expression (NPX) values from Olink were utilized. After further data processing, our study included a total of 52,632 individuals and 2,920 unique proteins (details see Supplementary Materials).

Genotype data were available for 502,493 participants in the UK Biobank v3 imputation. Detailed genotyping and quality control procedures performed by UK Biobank were described in a previous publication [46]. Our study excluded SNPs with call rates < 95%, minor allele frequency < 0.5% and deviation from the Hardy–Weinberg equilibrium with P < 1 × 10^−6^. Participants with less than 5% missing rates, not outliers in heterozygosity, who had no sex chromosome aneuploidy, of British ancestry, and who had no more than ten putative third-degree relatives in the kinship table were selected.

We used the nuclear magnetic resonance (NMR) metabolomics data in the UKB, which were recently released by Nightingale Health, containing around 292,000 individuals [47]. Here, a total of 249 metabolomics biomarkers were directly provided in the UKB and the details of these biomarkers could be found in Supplementary Table 13. More details for data processing were available in previous publications [47].

We used the brain structure imaging measures derived from T1 imaging data, as processed by WIN FMRIB on behalf of UKB [48]. The detailed preprocessing information was provided in Supplementary Materials. Both cerebral hemispheres and 66 regions defined by the Desikan-Killiany (DK) atlas for total CSA, total CV and mean CT were estimated [49].

### Calculation of *z*-scores

The general linear model was used to test the association between the proteomics/genomics/metabolomics data and brain-related traits. The effects of certain covariates (i.e. sex and age for plasma proteomics/metabolomics; sex, age and the first 20 genetic principal components for genomics) were regressed out. Then, we obtained an upper-tailed *p*-value. For each *p*-value, we performed a transformation based on the inverse normal cumulative distribution function (c.d.f.) into a *z*-score:

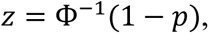

where Ф^−1^ was the inverse function of the standard normal cumulative distribution function.

### Mixture model-based replicability assessment (MMRA)

For two lists of z-scores: 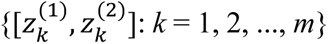, where *m* was the number of common units (i.e., proteins, SNPs and metabolites) from two different datasets, we considered a nine-component normal-mixture model for the joint distribution (see above for *z*-score calculation):

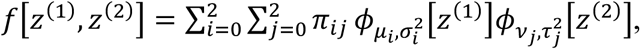

where 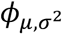 was the normal probability distribution function with mean 𝜇 and variance 𝜎^2^. We used the first component (index 0) to represent the null (no change/ association) feature component. Then, 𝜇_0_ = 𝜈_0_ = 0 and 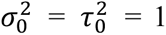. The second and third components (indices 1 and 2) were used to represent negative and positive associations. Their corresponding parameters (means and variances) were estimated from the paired *z*-scores with the following constraints: 𝜇_1_, 𝜈_1_ ≤ 0 and 𝜇_2_, 𝜈_2_ ≥ 0. 𝜋_𝑖𝑗_ was the proportion for component 𝑖 in the first association study and component 𝑗 in the second association study, and ∑_𝑖𝑗_ 𝜋_𝑖𝑗_ = 1.

This model was termed as partial concordance/discordance (PCD) model [17, 19]. Then, we defined 𝜋_𝑁𝑢𝑙𝑙_, 𝜋_*R*_ and 𝜋_*IR*_as the proportions of the non-signals, replicable, and irreplicable signals, respectively. These three latent categories were represented by:

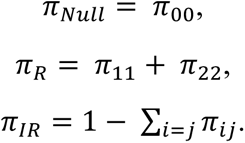

We reported an overall irreplicability quantity 𝜌_*IR*_ to measure the relative proportion of irreplicable signals in non-null signals:

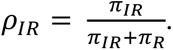

Additionally, for each unit *k* in an association study, we also defined the posterior probability of replicability as follows:

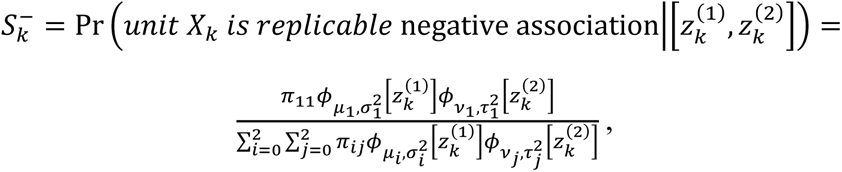

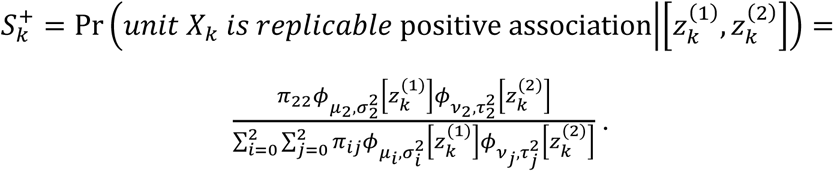

This estimated probability 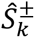 of 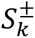 could be calculated by plugging-in the estimated parameters in the PCD model. Similarly, we also reported an individual irreplicability quantity 𝛾_𝑘_ to measure the relative proportion of irreplicable positive/negative association and no association for each unit *k* in an association study:

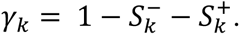

In this context, a lower 𝛾_𝑘_ value indicated a higher probability of replicable positive/negative association for unit *k*.

### Simulation design

Our simulations were designed based on the proteomics data from the UK Biobank. First, we split the original data into two subsets (referred to as Data 1 and Data 2 based on the order of subject number) with equal sample sizes. Then, we partitioned each subset randomly into two further subsets (referred to as Data 1A, Data 1B, Data 2A and Data 2B). Before the analysis, we ensured that sex, age was statistically similar between Data 1A versus 2A as well as Data 1B versus 2B (*t* test for age and chi-square test for sex, *P* > 0.05). Otherwise, we repeated the random data partition until this similarity requirement was satisfied. For each feature, there was no statistically significant differences in distribution between Data 1A versus 2A nor Data 1B versus 2B.

To generate upward or downward changes, a protein set was randomly chosen and an adjustment of 0.0123-0.0369 standard deviations of all the protein expression (corresponding to approximately 1-3 effect sizes in *z*-scores) were randomly added to or subtracted from the expression levels of the chosen protein set for each subject in Data 1A and Data 1B. This procedure was repeated 1,000 times. For each repetition, we obtained two lists of *z*-scores: one by protein-wisely comparing Data 1A versus Data 2A and the other by comparing Data 1B versus Data 2B. *Z*-scores were calculated based on the traditional two-sample *t*-test. A pair of *z*-scores were obtained for each protein. The replicability between two lists of *z*-scores was assessed by the MMRA approach. The following three simulations were considered.

a. *No un-replicable signal*. According to our random data partition, there were no statistically significant differences between Data 1A versus 2A or Data 1B versus 2B. We modified the 100% of null (no change) to 60% null, 20% upward changes and 20% downward changes as follows. We randomly selected two protein sets, each with 20% of the total proteins. To simulate 20% upward changes, for each protein in the first set, we randomly added a value equivalent to 1-3 effect sizes in *z*-scores to each subject’s protein expression levels in Data 1A and repeated this process in Data 1B to ensure 20% replicable upward changes. For each protein in the second set, we randomly subtracted a value equivalent to 1-3 effect sizes in *z*-scores from each subject’s protein expression in Data 1A and repeated this process in Data 1B to achieve 20% replicable downward changes.
b. *Moderate level of un-replicable signal*. We randomly selected four protein sets. The first two sets each comprised 15% of the total proteins, and the upward changes and downward changes were simulated as described in (a). The next two sets each comprised 5% of the total proteins. For each protein in the third set, we randomly added a value equivalent to 1-3 effect sizes in *z*-scores to each subject’s protein expression level in Data 1A (but not in Data 1B). Then, we had 5% discordant changes (up versus null). Similarly, for each protein in the fourth set, we subtracted a value from each subject’s protein expression level in Data 1A (but not in Data 1B) so that we had 5% discordant changes (down versus null).
c. *High level of un-replicable signal*. Considering that the replicability levels may vary across different studies, we randomly selected four protein sets, each with 10% of the total proteins. The replicable upward/downward changes (the first/second set) and un-replicable upward/downward changes (the third/fourth set) were simulated similarly as described in (b).

### Overall replicability assessment

To investigate the overall replicability (𝜌_*IR*_) for proteomics-based association study (namely PBAS) results with different phenotypes, we considered a random subsampling approach which randomly split the whole data into two subsets with (approximately) equal sample sizes for 1,000 times. Due to constraints from missing observations, different sample sizes were included in different PBAS scenarios. Aside from proteomics data, we used the following four types of measures in our study: physical measures, cognitive function, mental health and brain imaging metrics (details for each trait see Supplementary Table 1). The brain structure imaging metrics included total cortical surface area (CSA), total cortical volume (CV) and mean cortical thickness (CT). We calculated the overall replicability ( 𝜌_*IR*_) for these brain metrics in both cerebral hemispheres (*N* = 5,623). We also assess the overall replicability for these metrics based on the Desikan–Killiany (DK) atlas which included 66 regions. Using the above data, we obtained 1,000 pairs of *z*-scores and assessed the median 𝜌_*IR*_ for each PBAS scenario.

### Effects of season and fasting time at blood collection

To assess the effects of season and fasting time on replicability, we further included these factors as additional covariates in the PBAS. The season of blood collection was categorized as summer/autumn (June to November) or winter/spring (December to May), based on the blood collection date. Fasting time was determined from participant-reported fasting duration prior to blood collection.

### Multi-omics comparison

The protein quantitative trait locus (pQTL) identified in a previous study [1] were adopted to compare the overall replicability level between genetics-based association analyses and proteomics-based association analyses. After quality control, 43,685 participants with both genetics and proteomics data were included for this analysis. Each participant had 2,920 proteins and their corresponding 6,386 pQTL-related SNPs in UKB [1]. Moreover, to compare the overall replicability level between metabolomics-based association analyses and proteomics-based association analyses, we then included 30,079 participants with both metabolomics and proteomics data. The PBAS was conducted based on the same procedure as mentioned in the previous section. The same procedure was also employed to analyze the associations between the genetics/metabolomics data and each brain-related measure (within physical measures, cognitive function, mental health and brain imaging metrics). Then, we obtained 1,000 pairs of z-scores based on 1,000 random subsampling times; and we calculated the median 𝜌_*IR*_ for each association analysis scenario.

The PBAS scenarios considered here excluded subjects without British ancestry (see Study Participants Section in Methods for details) when compare the overall replicability level between genetics-based association analyses and proteomics-based association analyses.

### Influential factors

According to the details of UKB proteomics data collection (http://biobank.ndph.ox.ac.uk/ukb/ukb/docs/Olink_proteomics_data.pdf), two versions of the assay panel were employed by UKB to collect proteomics data which included “Explore 1,536 assay panels” and “Expansion 1460 assay panels” (referred as Panel 1 and Panel 2, respectively).

Here, we stratified all plasma proteins into three categories based on their dilution levels: 1,941 proteins at least-abundant (1:1), 524 proteins at moderate-abundant (1:10) and 455 proteins at more-abundant (1:100 to 1:100,000). Moreover, we further stratified the proteins into four categories according to the proportion of samples below the LOD: <0.05% (1,012 proteins), 0.05%-1% (707 proteins), 1%-50% (544 proteins) and >50% (657 proteins). Then, we counted the number of proteins across twelve categories (i.e., three dilution level categories × four proportion of samples below the LOD categories) in Panel 1 and Panel 2. To evaluate the impact of sample size on overall replicability, we randomly selected two settings with approximately 50% and 25% sample size from the original dataset. Then, we split each setting into two subsets with (approximately) equal sample sizes. This process was repeated 1,000 times. We then calculated the overall irreplicability quantity 𝜌_*IR*_based on each of these subsets.

### Prediction modeling

The proteomics technologies are currently under active developments, and more proteomics panels to include additional proteins may be made available in the near future. Here, we assumed that a total of 1,460 proteins would still be included in a future proteomics panel, and we intended to estimate the related overall replicability of associations (based on simulations). For this analysis, given a phenotype and a simulated panel, we defined the term “overall replicable” as 𝜌_*IR*_ < 0.05 (i.e., binary response variable). With the dilution level (divided into three categories), the proportion of samples below the LOD (divided into four categories), and the sample size, we can construct a logistic regression model for this purpose. Please see Supplementary Materials for the related details and comprehensive simulation results.

### Individual replicability assessment

To assess the individual replicability (𝛾_𝑘_) of proteins in the PBAS within cognitive function and mental health, we conducted further analyses for each PBAS scenario which exhibited high levels of overall replicability (median overall irreplicability quantity 𝜌_*IR*_ < 0.05). Initially, we calculated the median individual irreplicability quantity 𝛾_𝑘_ for each protein in 1,000 random subsampling times. These median 𝛾_𝑘_ values were then ranked in an ascending order.

In this study, we identified the pleotropic proteins for cognitive function and mental health that demonstrated the highest replicability associations with more than 50% of the traits under investigation. The pleotropic proteins were selected based on the following criteria: they must be identified as one of the top ten proteins in at least 10 mental health-related traits (more than 50% of total 19 traits) or 3 cognitive function-related traits (more than 50% of total 5 traits). The random subsampling approach (See *Overall replicability assessment* section in Methods) was used to simulate practical study cohorts and their replications. For each random subsampling, we defined individual irreplicability quantity 𝛾_𝑘_ < 0.05 as the criterion to indicate that a protein exhibited directional consistency. For each trait in the cognitive function and mental health, the fraction of directional consistency (FDC) was defined as the fraction of times that a protein demonstrated directional consistency (𝛾_𝑘_ < 0.05) across 1,000 random subsampling.

We further investigated the impact of proteomics data quality on individual replicability for proteins. Here, for each trait under cognitive function and mental health with median overall irreplicability quantity 𝜌_*IR*_ < 0.05, we calculated median individual irreplicability quantity 𝛾_𝑘_ for each protein from 1,000 random subsampling times. Then, for all proteins’ median 𝛾_𝑘_, we reported the median value for Panel 1 versus Panel 2. We also analyzed the impact of dilution levels and the proportion of samples below the LOD for individual replicability performance.

## Data availability

The data used in the study from the UKB was accessible under restricted access (application number 19542). Access can be obtained by submitting an application through the UKB platform (https://www.ukbiobank.ac.uk/).

## Code availability

Code for overall/individual replicability assessment and predictive framework for potential future panels is openly shared in GitHub (https://github.com/YixinZhang-stat/plasma-proteomics-replicability). R version 4.3.0 and R package ggseg was used to show median 𝜌_*IR*_for each region in DK atlas.

## Acknowledgements

This work was partially supported by a start-up fund from the University of Science and Technology of China (to Yinglei Lai). National Key R&D Program of China (2019YFA0709502, 2018YFC1312904 to Jianfeng Feng), Shanghai Municipal Science and Technology Major Project (2018SHZDZX01 to Jianfeng Feng), 111 Project (B18015 to Jianfeng Feng), Humboldt Research Award (to Jianfeng Feng). Some image materials in Figure 1 were free acquired from Freepik (https://www.freepik.com).

## Author contributions

Conception: Zeyu Jiao, Yinglei Lai, Yixin Zhang and Jianfeng Feng. Design: Zeyu Jiao, Yixin Zhang, Yinglei Lai, Jujiao Kang and Jianfeng Feng. Data acquisition, processing and analysis: Zeyu Jiao, Yixin Zhang, Yinglei Lai, Jujiao Kang, Wei Zhao, Jia You, Wei Cheng and Jianfeng Feng. Manuscript writing and revising: Zeyu Jiao, Yixin Zhang, Yinglei Lai, Jujiao Kang, Liang Ma and Jianfeng Feng.

## Competing interests

The authors declare no competing interests.

## Notes

### Competing Interest Statement

The authors have declared no competing interest.

