## Supplementary Materials for "Merits and challenges of plasma proteomics on association replicability"

**Plasma proteomics**

The UKB resource is a large, prospective, and open-access biomedical database launched between 2006 and 2010 [1]. As a public-private partnership, the UKB cohort provides the scientific community with a considerable breadth and depth of health-related information. Over 500,000 eligible participants aged 40-69 years at baseline were enrolled from 22 assessment centers across the United Kingdom (England, Wales, and Scotland), with ongoing follow-up. Each participant completed a touch-screen questionnaire, a nurse-led interview, and an anthropometric examination. Detailed data on genotyping, imaging, electronic health record linkage, biospecimens, and body measurements were collected. In this study, we used the phenotypic data from the initial assessment visit (2006-2010), during which participants were recruited and consented.

The Pharma Proteomics Project was a precompetitive biopharmaceutical consortium that characterized the plasma proteomic profiles of 55,323 UKB participants. As outlined in the Supplementary Information of previous article [2], proteomic data underwent stringent quality control procedures. The proteins levels were provided as Normalized Protein eXpression (NPX) values after log transformation and normalization within and across batches. The UK Biobank Pharma Proteomics Project further removed outliers and data labeled with QC warnings suggested by Olink's facilities. No additional NPX processing was performed in this study after these quality control measures.

Among the 55,323 participants with proteomic data, measurements were taken for 2,923 unique proteins. A total of 53,026 participants who had baseline measurements were included in the analysis. Participants missing over 60% of the protein data and proteins missing in over 60% of participants were excluded [3]. This resulted in a final dataset consisting of 52,632 individuals and 2,920 proteins. Missing data were imputed using the median values within each gender group and age category (0-25%, 25-50%, 50-75%, and 75-100% age percentiles) [3].

**MRI acquisition and preprocessing**

Magnetic resonance imaging (MRI) of the UK biobank was performed using a

Siemens Skyra 3T running VD13ASP4 (Siemens Healthcare, Erlangen, Germany) with a Siemens 32-channel RF receive head coil. The T1 structural protocol was acquired at 1 mm isotropic resolution using a three-dimensional (3D) MPRAGE acquisition, with inversion and repetition times optimized for maximal contrast. The superior inferior field-of-view (FoV) was large (256 mm), at little cost, to include reasonable amounts of neck/mouth, which might be of interest to some researchers (for example, in the study of sleep apnea).

For raw T1 imaging data, the FoV was reduced to minimize the amount of non-brain tissue and gradient distortion correction was applied. The reduced-FoV T1 imaging data were nonlinearly warped to MNI152 space using FNIRT [4]. Next, the T1 images were processed with FreeSurfer [5]. The primary FreeSurfer modelling focused on the cortical surface. Surface atlases were used to extract IDPs relating to standard atlas regions' surface area, volume and mean cortical thickness.

### Simulation results for the *MMRA* approach

The simulation results were summarized in Figure S1. In simulation (a) with no un-replicable signals, the median values of $\pi_R$, $\pi_{IR}$ and $\rho_{IR}$ were 0.3784, 0.0101 and 0.0277, respectively. Furthermore, the related lower- and upper-quartiles (Q1-Q3) were 0.3608-0.4124, 0.0003-0.0475 and 0.0008-0.1123, respectively. Given the lower-quartiles of $\pi_{IR}$ and $\rho_{IR}$ were close to 0, it was reasonable to conclude that the assessed results showed no un-replicable signals in this simulation scenario. In simulation (b) with a moderate level of un-replicable signals (25%), the median values of $\pi_R$, $\pi_{IR}$ and $\rho_{IR}$ were 0.3030, 0.0836 and 0.2139 while the related Q1-Q3 were 0.2760-0.3586, 0.0532-0.1383 and 0.1397-0.3078, respectively. In simulation (c) with high level of un-replicable signals (50%), the median values of $\pi_R$, $\pi_{IR}$ and $\rho_{IR}$ reached 0.2423, 0.1661 and 0.5906 with the Q1-Q3: 0.1918-0.3175, 0.1206-0.2326 and 0.2982-0.5165, respectively. For simulation (b) and (c), we could observe the true values of $\pi_R$, $\pi_{IR}$ and $\rho_{IR}$ consistently fell within the range of Q1-Q3. These simulation findings demonstrated that the MMRA approach could accurately assess the replicability of PBAS results.

**Replicability differences between brain imaging measures**

To explore the potential explanations for the observed differences in overall replicability levels between total CSA/CV and mean CT, we calculated the one-sided p-values and z-scores from PBAS based on the entire data for total CSA, total CV and mean CT in both hemispheres. Then, we estimated the proportion of true null hypotheses ($\pi_0$) based on p-values by using "convest" function from limma package (https://www.rdocumentation.org/packages/limma/versions/3.28.14/). For total CSA and CV, the $\pi_0$ values were consistently around 0.65 in both cerebral hemispheres. In contrast, mean CT showed relatively higher $\pi_0$ values, reaching 0.760 in the left hemisphere and 0.784 in the right hemisphere (Figure S4a, b). Specifically, the histograms of p-values for total CSA, total CV in the both hemispheres exhibited a clear bimodal distribution, with peaks near p-values of 0 and 1. This pattern suggested the presence of positive/negative significant associations (one-sided $p$-values close to 0 and 1). However, the histograms of $p$-values for mean CT demonstrated a less pronounced bimodal feature, with broader distributions of $p$-values. This wider spread indicated fewer positive/negative significant associations in the results.

We further evaluated the z-scores for different brain structure metrics (Figure S4c). The z-scores for mean CT close to a standard normal distribution, whereas z-scores for total CSA and CV shown heavy-tailed characteristics, indicating stronger association strengths. Besides, we assessed the concordance of $z$-scores for different brain structure metrics between the left and right hemispheres. The scatter plots of $z$-scores also displayed clearer alignment between hemispheres for total CSA and total CV compared to mean CT. Total CSA exhibited a strong concordance, with a Pearson correlation coefficient of $r =$ 0.997, and a similarly high concordance was observed for total CV ($r =$ 0.979). In contrast, mean CT demonstrated a clear reduction in concordance between hemispheres, with $r =$ 0.869. In conclusion, these findings indicated that mean CT exhibited relatively weaker association strength than total CSA and CV in the PBAS.

**Replicability prediction procedures for potential future panels**

To generate data for training the model (with adequate variations based on the current two panels), our protein sampling procedure was designed as follows. A total of 1460 proteins were randomly sampled from both panels using a binomial mixture. Specifically, $r \sim Binomial(1460, p)$ proteins were sampled from Panel 1, and the remaining from Panel 2, with $p = i/101,\ i = 1, \ldots, 100.$ This yielded 1,000 synthetic protein panels with controlled proportions. Furthermore, For each synthetic panel, a random sample size $n \in [0.6N, N]$ (where $N$ is the total number of subjects) was drawn. Additionally, we constructed 24 controlled scenarios, each corresponding to one specific combination (dilution level × proportion of samples below LOD × Panel). For each scenario, 1,000 synthetic panels were generated such that a target subset of proteins (based on dilution level/proportion of samples below the LOD/Panel combinations) were excluded from the sampling set. This yielded $3 \times 4 \times 2 \times 1000$ =24,000 synthetic protein panels. Here each synthetic instance included nine variables, the calculated irreplaceability quantity $\rho_{IR}$, the sample size used in the association test, and the counts of proteins in each of seven predefined categories based on dilution levels and proportion of samples below the LOD categories. These categories were denoted as DL1 (1:1), DL2 (1:10), DL3 (>1:10) for dilution levels and PLOD1 (<0.05%), PLOD2 (0.05%-1%), PLOD3 (1%-50%), PLOD4 (>50%) for LOD-based categories. Repeat above procedure two times, we had 50,000 synthetic protein panels. Then we split them into train and test datasets with a ratio of 1:1, each with 25,000 synthetic protein panels.

Based on the scatter plots of these variables against quantile normalized $\rho_{IR}$ (Figure S9), the DL1, PLOD1 and sample size were selected as predictors for the subsequent simplified model as they captured the most distinct and directionally consistent associations with replicability. Specifically, among the three dilution-level categories (DL1, DL2, and DL3), DL1 showed a positive correlation with quantile-normalized $\rho_{IR}$, while DL2 and DL3 were negatively correlated. In contrast, among the LOD-based categories, PLOD1 exhibited a strong negative correlation, whereas PLOD2–PLOD4 showed weak or positive correlations. Notably, sample size was also negatively correlated with quantile-normalized $\rho_{IR}$, supporting the need to include this variable in model construction. We then binarized the irreplaceability quantity $\rho_{IR}$ with threshold 0.05 and adopted the logistic regression model to estimate the related overall replicability of associations (based on simulations). We fit standard logistic regression models on the train set to predict the replicability outcome. For traits like fluid intelligence and neuroticism, The AUC-ROC curve and Calibration curve based on test set were displayed in Figure S10. The logistic regression models were specified as:

$$logit\left(\mathbb{I}_{\rho_{IR}<0.05}\right) = intercept_{traits} + DL1 + PLOD1 + \log_{10}(sample\ size).$$

To interpret how different combinations of variables influence the overall replicability, we designed four representative simulation scenarios. Panel 1: empirical distributions of dilution level and proportion of samples below the LOD based on UKB Panel 1; Panel 2: empirical distributions based on UKB Panel 2; Scenario1: DL1 = 0, PLOD1 = 1 (high quality); Scenario 2: DL1 = 1, PLOD4=1 (low quality). Each structure was repeated at 10 sample sizes (10,000 to 100,000 in increments of 10,000), yielding 40 synthetic proteomics panels. Predictions from the selected final model showed clear separation: The high-quality scenario (Scenario 1) consistently achieved the highest replicability probability, while the low-quality scenario (Scenario 2) remained low across all sample sizes. Panel 1 and Panel 2 scenarios revealed intermediate levels consistent with their respective data characteristics. Notably, the effect of sample size varied across traits. For traits such as fluid intelligence, predicted replicability probability increased rapidly with larger sample sizes (Figure S10 a). However, for traits like neuroticism, increasing sample size yielded moderate improvement in replicability (Figure S10 c). These results indicated that the benefit of larger sample size varied across traits. Moreover, we also fit the inverse probability–weighted (IPW) logistic regression models to predict the replicability outcome [6]. Similar trends could be observed in our results (Figure S10 b, d).

**Individual replicability assessment for Cardiometabolic, Inflammation, Neurology and Oncology panels**

According to the details of UKB proteomics data collection (http://biobank.ndph.ox.ac.uk/ukb/ukb/docs/Olink_proteomics_data.pdf), the "Explore 1,536 assay panels" included a range of sub-panels, namely Cardiometabolic, Inflammation, Neurology and Oncology. Similarly, the "Expansion 1,460 assay panels" included Cardiometabolic II, Inflammation II, Neurology II and Oncology II. Here, for each phenotype within cognitive function and mental health, we calculated median $\gamma_k$ for each protein in Cardiometabolic vs. Cardiometabolic II (366 and 366 proteins, respectively), Inflammation vs. Inflammation II (365 and 365 proteins, respectively), Neurology vs. Neurology II (363 and 364 proteins, respectively) and Oncology vs. Oncology II (367 and 364 proteins, respectively). Then, for all proteins' median $\gamma_k$, we reported the median value for each panel. For all the phenotypes within cognitive function and mental health, the irreplicability level for Cardiometabolic, Neurology and Oncology was clearly higher compared to the irreplicability level for Cardiometabolic II, Neurology II and Oncology II, respectively (Figure S11 a, b, c). For Inflammation vs. Inflammation II, we also observed higher irreplicability level for Inflammation in some traits within cognitive function and mental health ((Figure S11 b)). In other traits, the irreplicability level was similar between Inflammation and Inflammation II.

For each assay panels, we calculated the data missing rate for each protein. For Cardiometabolic vs. Cardiometabolic II, the median data missing rate was 0.0270 and 0.1717 while Q1-Q3 was 0.0235-0.0434 and 0.1494-0.1770, respectively. For Inflammation vs. Inflammation II, the median data missing rate was 0.0263 and 0.1691 while Q1-Q3 was 0.0215-0.0296 and 0.1579-0.1749, respectively. For Neurology vs. Neurology II, the median data missing rate was 0.0312 and 0.1726 while Q1-Q3 was 0.0242-0.0456 and 0.1508-0.1726, respectively. For Oncology vs. Oncology II, the median data missing rate was 0.0280 and 0.1648 while Q1-Q3 was 0.0213-0.0456 and 0.1533-0.1798, respectively.

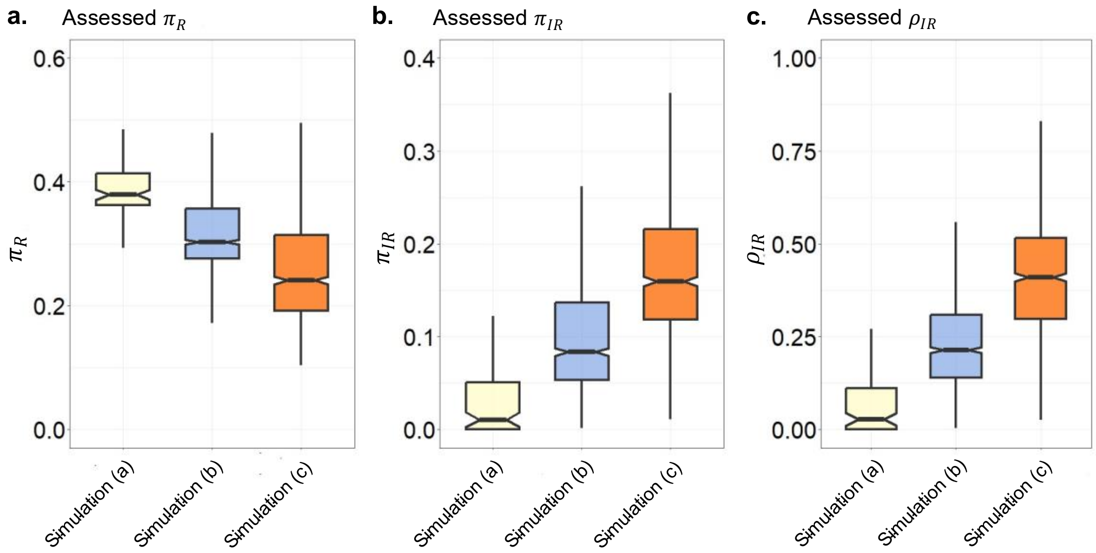


**Figure S1| Simulation-based assessment of overall replicability. a-c,** Box plots of assessed $\pi_R$, $\pi_{IR}$ or $\rho_{IR}$ across three simulation designs.

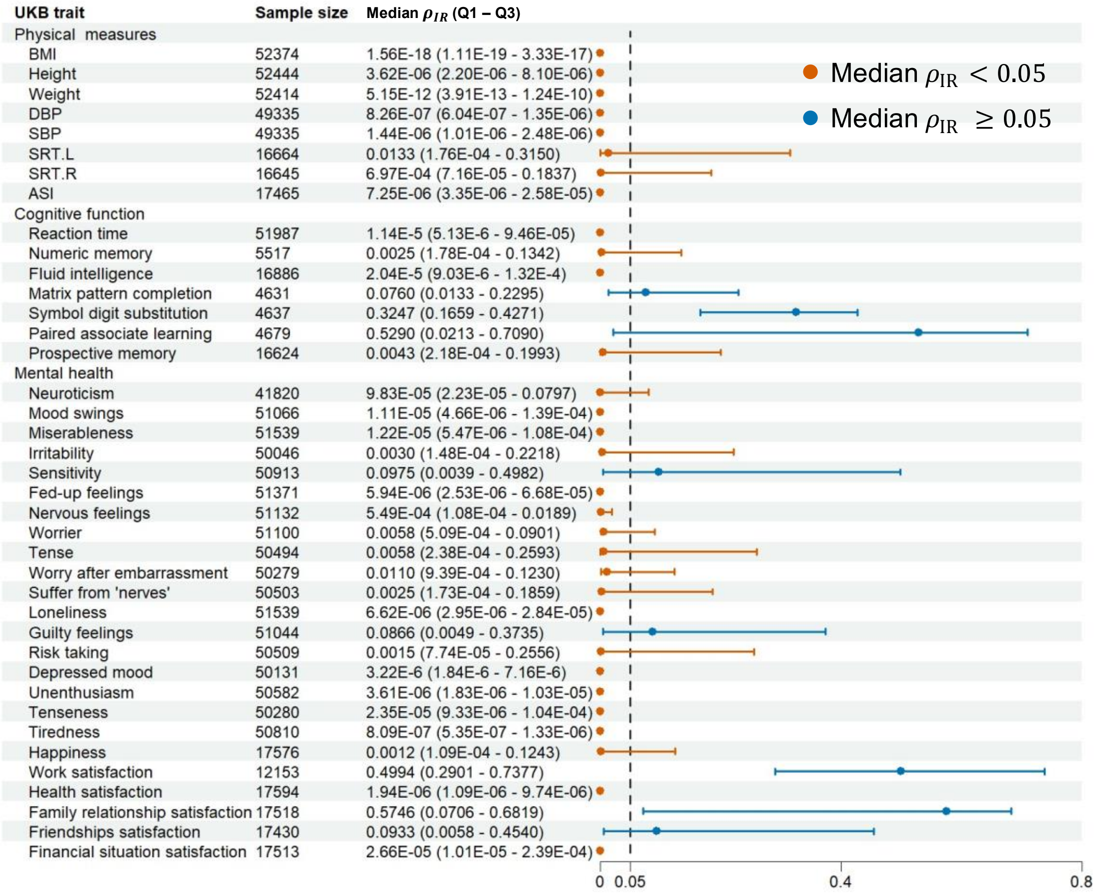


**Figure S2 | Overall replicability assessment of PBAS adjusted for additional covariates.** PBAS were additionally adjusted for season and fasting time. For each trait, sample sizes were indicated, and the median overall irreplicability quantity $\rho_{IR}$ together with lower- and upper-quartiles (Q1-Q3) were estimated from 1,000 times random subsampling. Orange points denoted traits with median $\rho_{IR} < 0.05$, and blue points denoted traits with median $\rho_{IR} \geq 0.05$.

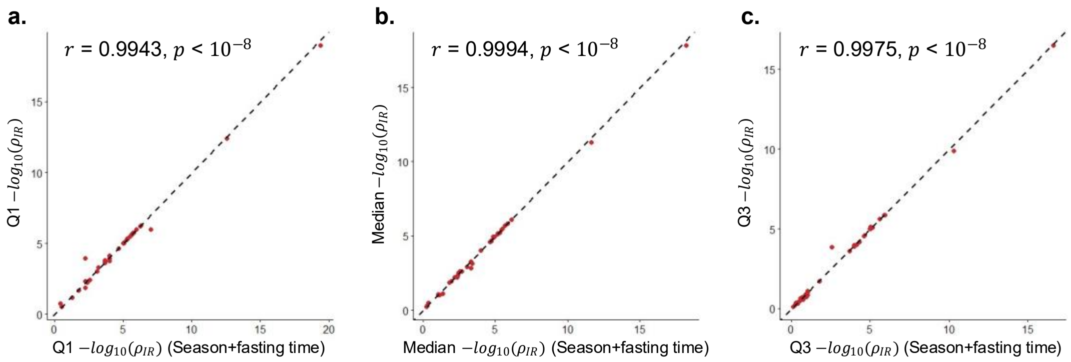


**Figure S3 | Consistency of overall replicability after covariate adjustment.** Scatter plots of (**a**) the lower quartile Q1, (**b**) median, and (**c**) upper quartile Q3 of $-\log_{10}(\rho_{IR})$ before versus after adjustment for season and fasting time. Each point represented a trait. The dashed line denoted the diagonal. Pearson correlation coefficients (*r*) and corresponding *p*-values were reported.

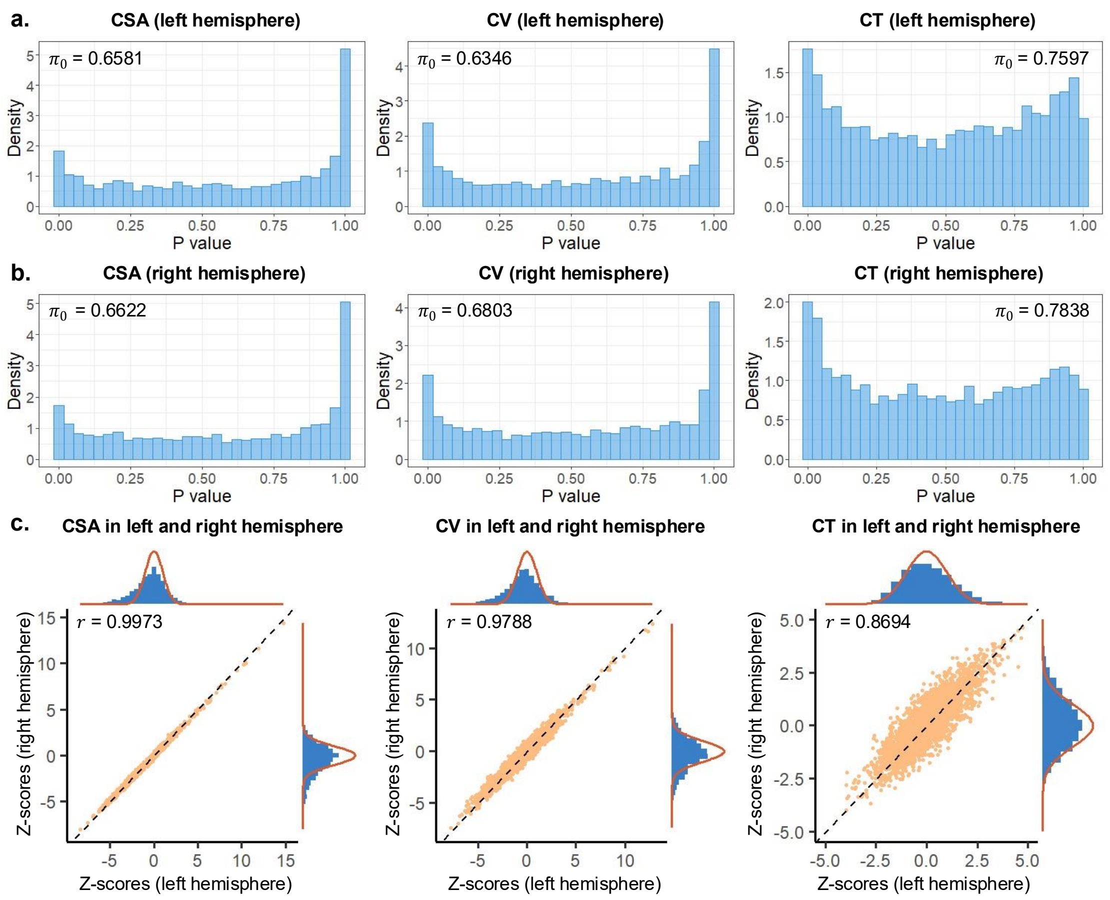


**Figure S4 | Distribution of association statistics for brain imaging metrics. a,** Histograms of $p$-values from PBAS for total CSA, total CV, and mean CT in the left hemisphere. $\pi_0$ represented the estimated proportion of true null hypotheses. **b,** Histograms of $p$-values for total CSA, total CV, and mean CT in the right hemisphere. **c,** Scatter plots of $z$-scores for total CSA, total CV, and mean CT between the left and right hemispheres. Each point represents a protein - trait association. The dashed line denoted the diagonal, and red curves indicated the density of standard normal distribution.

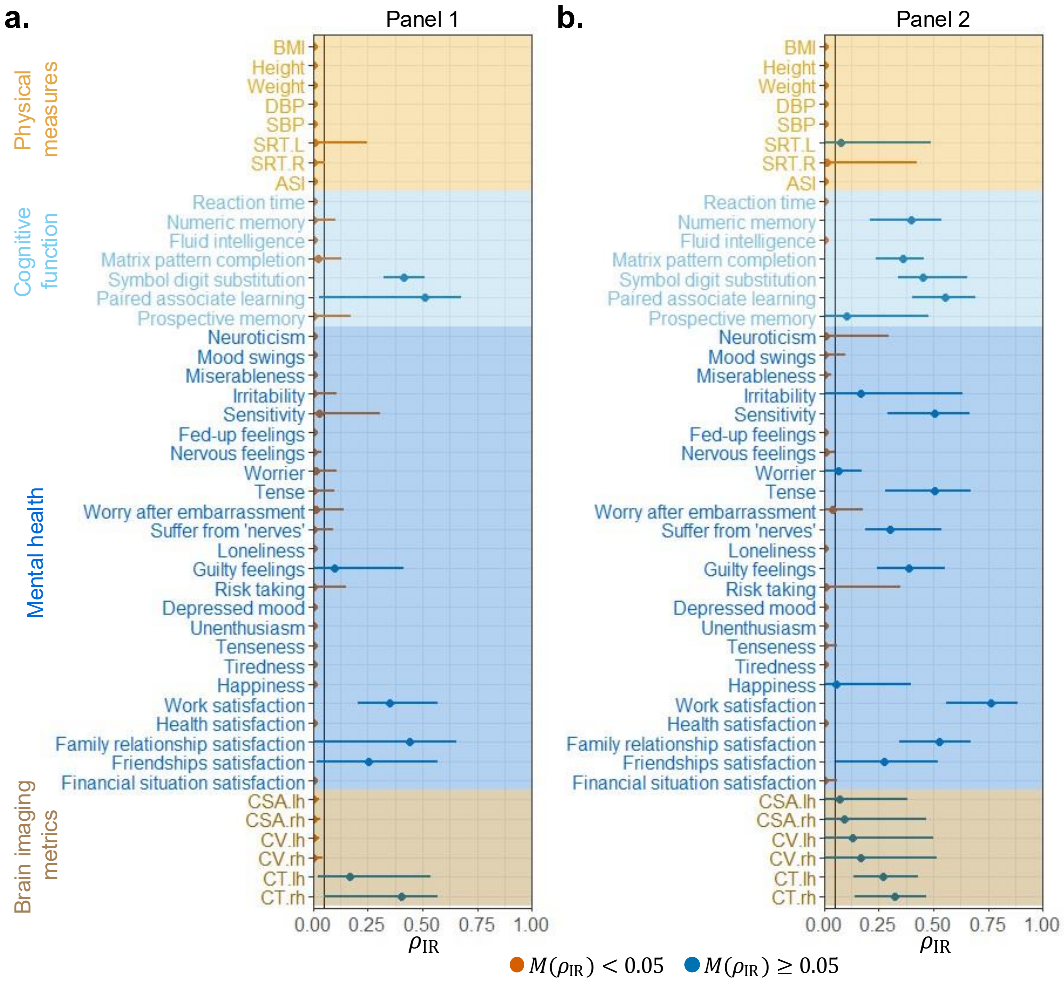


**Figure S5 | Comparison of overall replicability between two proteomic panels. a,** Estimated overall irreplicability quantity $\rho_{IR}$ for Panel 1 across traits. **b,** Corresponding estimates for Panel 2. For each trait, the median $\rho_{IR}$ with lower- and upper- quantiles were obtained from 1,000 random subsampling times. Orange points denoted traits with median $\rho_{IR} < 0.05$, and blue points denoted traits with median $\rho_{IR} \geq 0.05$.

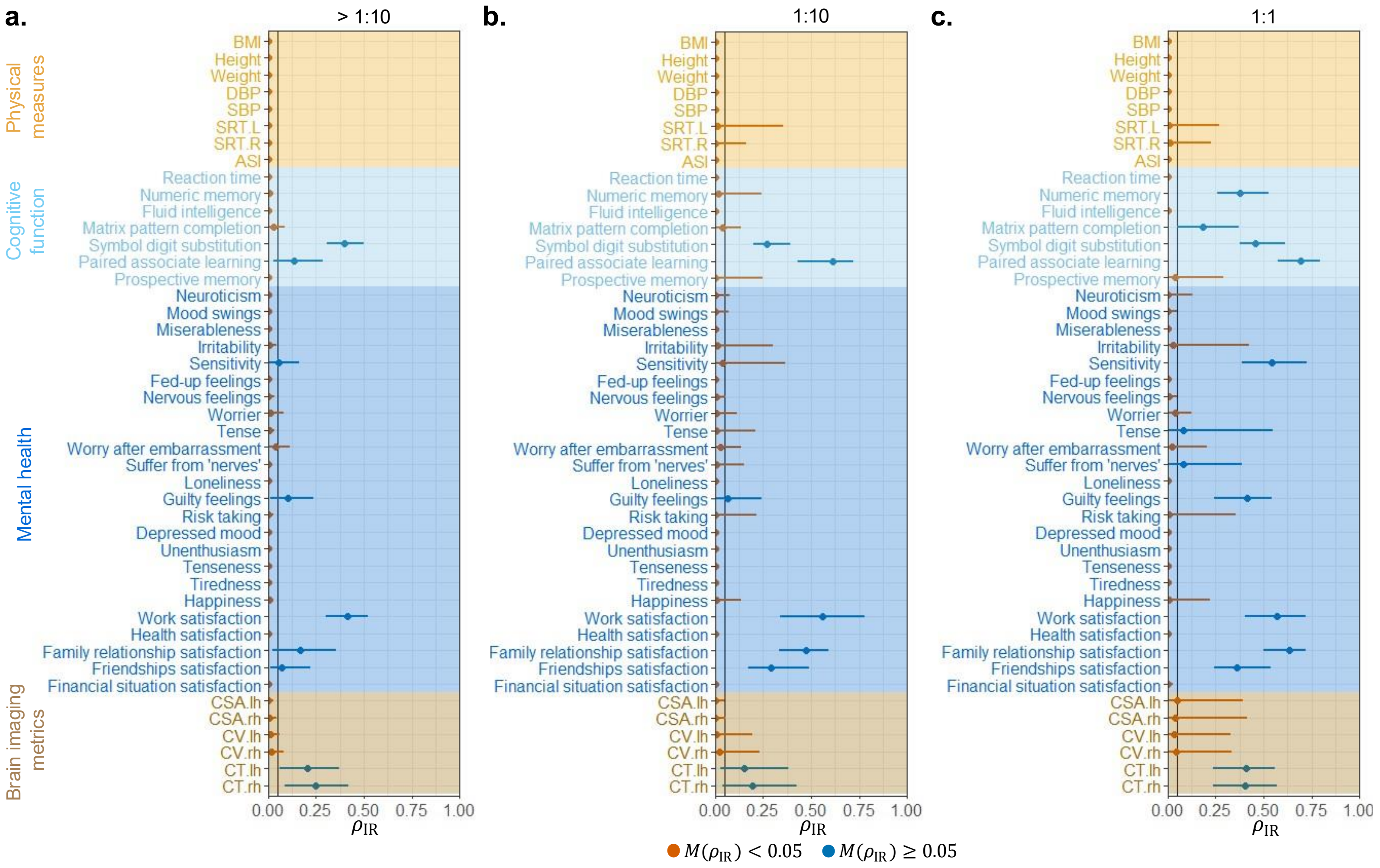


**Figure S6 | Impact of dilution levels on overall replicability.** Estimated overall irreplicability quantity $\rho_{IR}$ across traits for PBAS under different proteomics dilution levels: (**a**) >1:10, (**b**) 1:10, and (**c**) 1:1. For each trait, the median $\rho_{IR}$ with lower- and upper- quantiles were obtained from 1,000 random subsampling times. Orange points denoted traits with median $\rho_{IR} < 0.05$, and blue points denoted traits with median $\rho_{IR} \geq 0.05$.

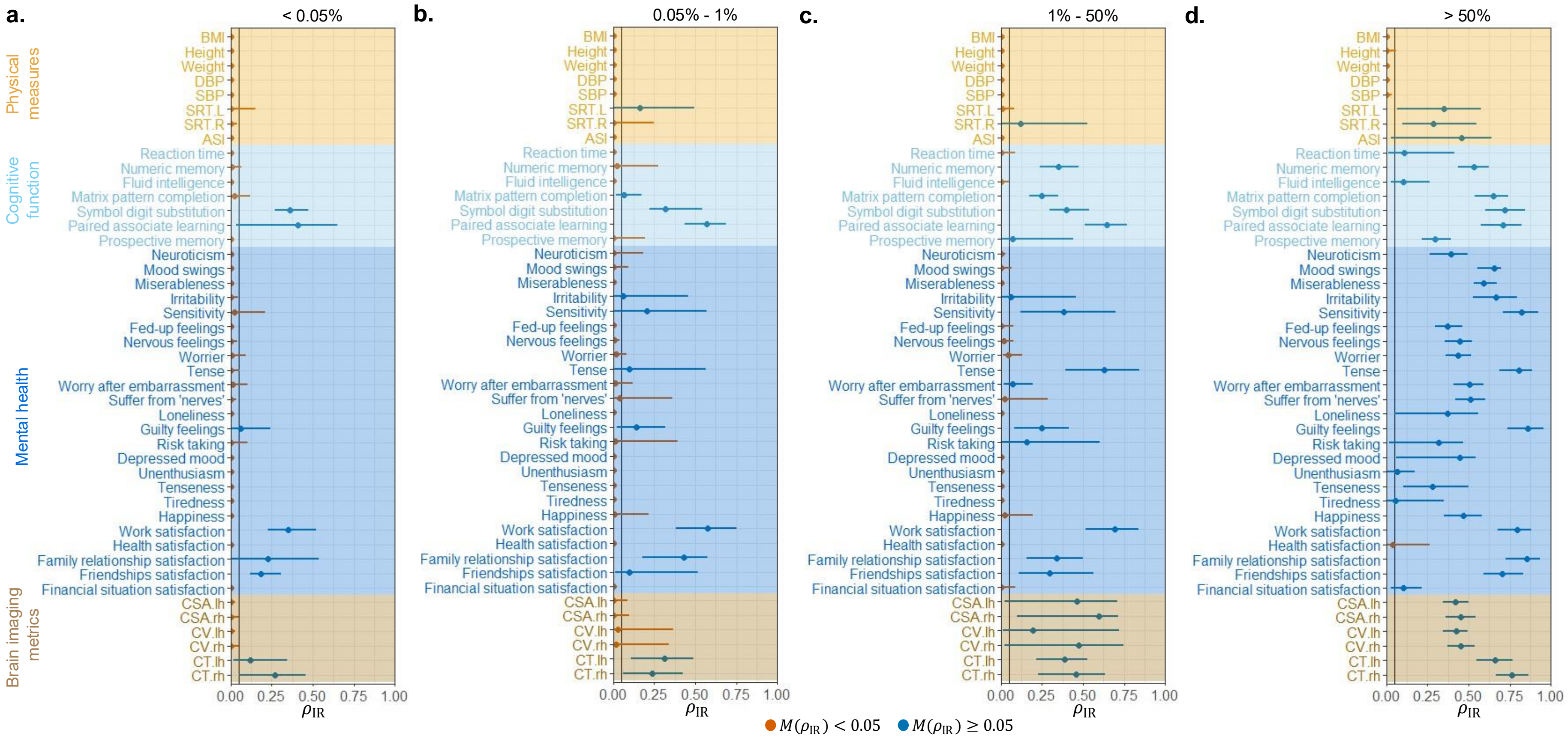


**Figure S7 | Impact of the proportion of samples below the LOD on overall replicability.** Estimated overall irreplicability quantity $\rho_{IR}$ across traits for PBAS under different proportions of samples below the LOD: (**a**) <0.05%, (**b**) 0.05–1%, (**c**) 1–50%, and (**d**) >50%. For each trait, the median $\rho_{IR}$ with lower- and upper- quantiles were obtained from 1,000 random subsampling times. Orange points denoted traits with median $\rho_{IR} < 0.05$, and blue points denoted traits with median $\rho_{IR} \geq 0.05$.

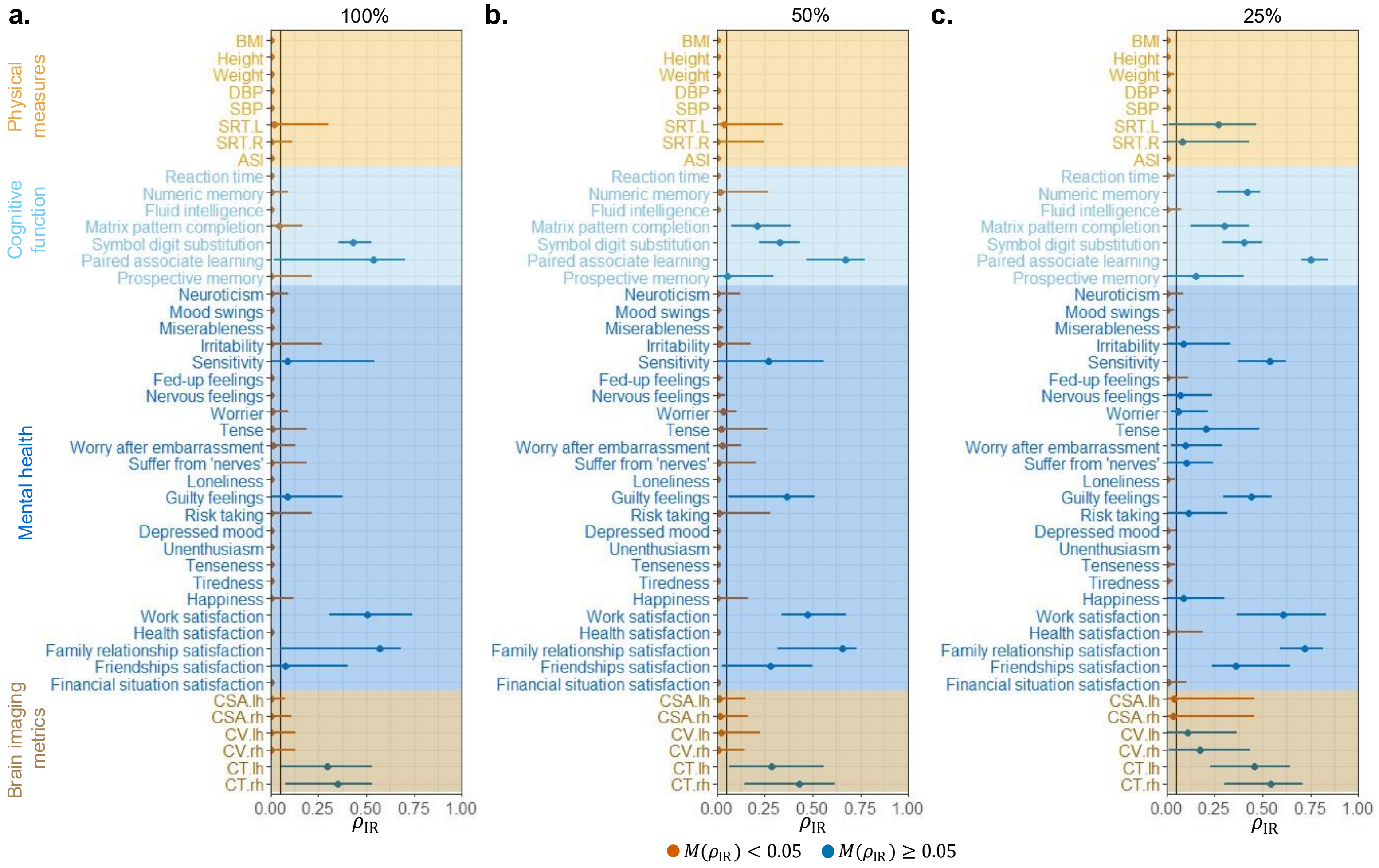


**Figure S8 | Impact of sample size on overall replicability.** Estimated overall irreplicability quantity $\rho_{IR}$ across traits across traits for PBAS under different sample sizes: (**a**) 100%, (**b**) 50%, and (**c**) 25% of whole data. For each trait, the median $\rho_{IR}$ with lower- and upper- quantiles were obtained from 1,000 random subsampling times. Orange points denoted traits with median $\rho_{IR} < 0.05$, and blue points denoted traits with median $\rho_{IR} \geq 0.05$.

**a.** Fluid Intelligence

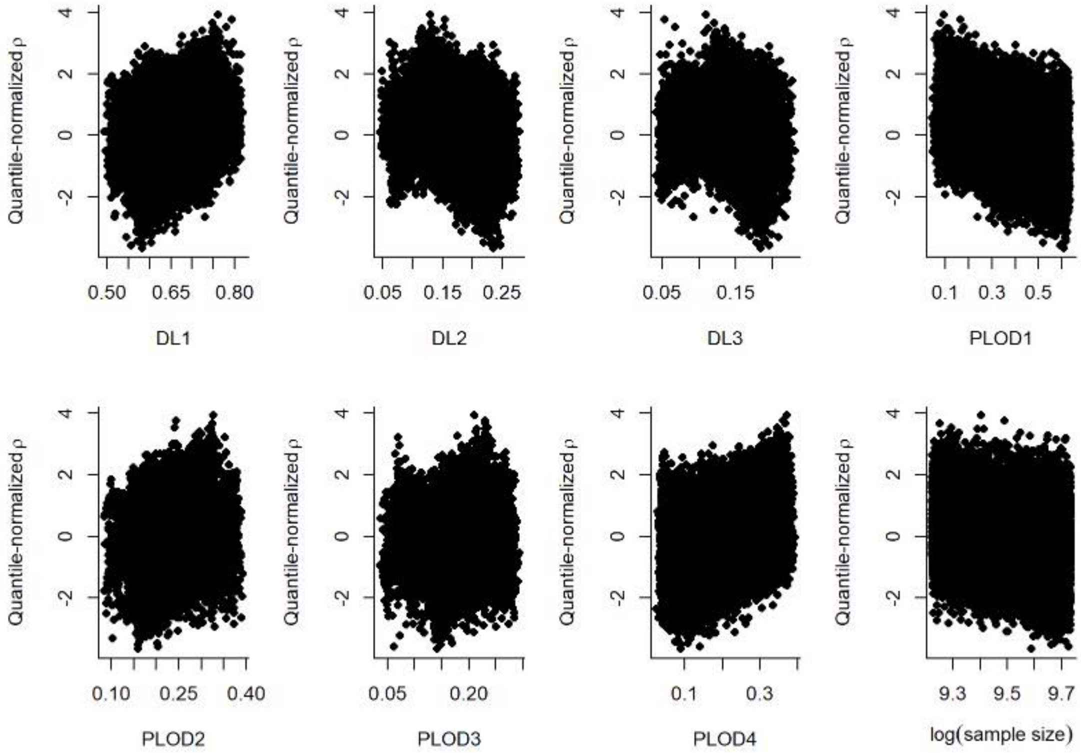


**b.** Neuroticism

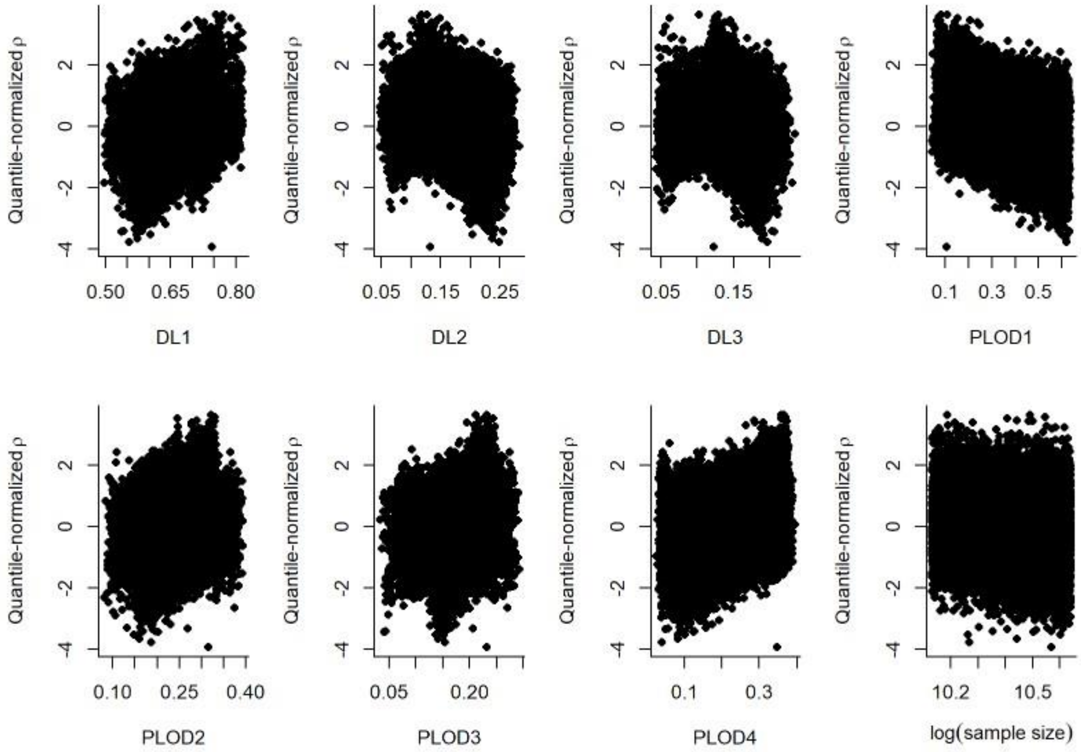


**Figure S9 | Associations of seven predefined categories and sample size with quantile-normalized overall replicability.** Scatter plots of relationship between quantile-normalized overall replicability and seven predefined categories, together with sample size. **a**, Results for fluid intelligence. **b**, Results for neuroticism. Categories include dilution levels (DL1-DL3), proportions of samples below the LOD (PLOD1-PLOD4), and log-transformed sample size.

**a.** Fluid Intelligence (standard)

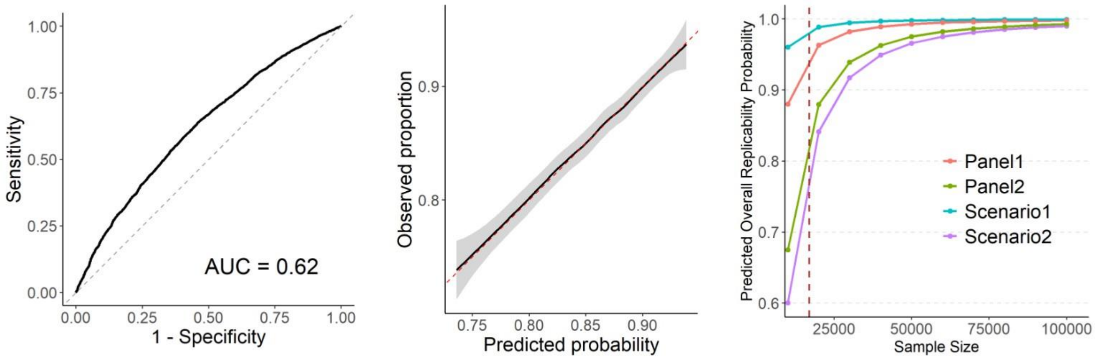


**b.** Neuroticism (standard)

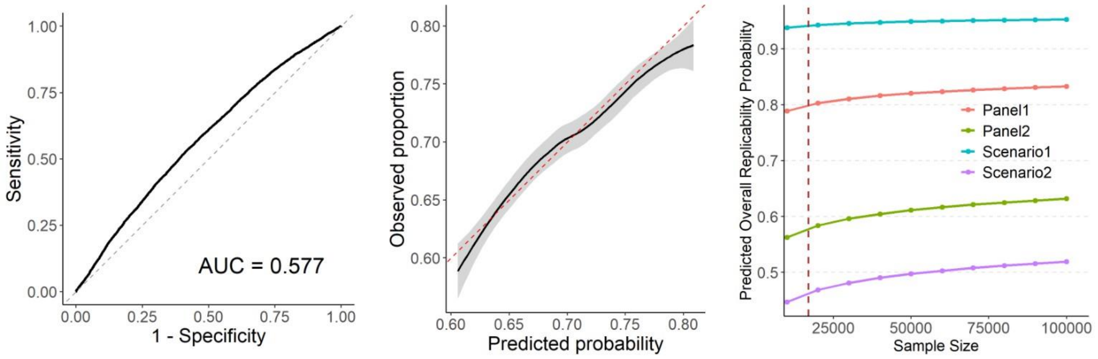


**c.** Fluid Intelligence (IPW)

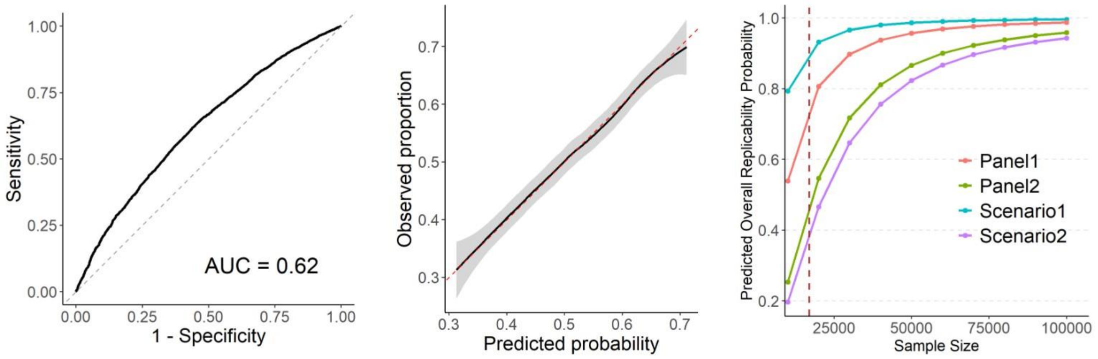


**d.** Neuroticism (IPW)

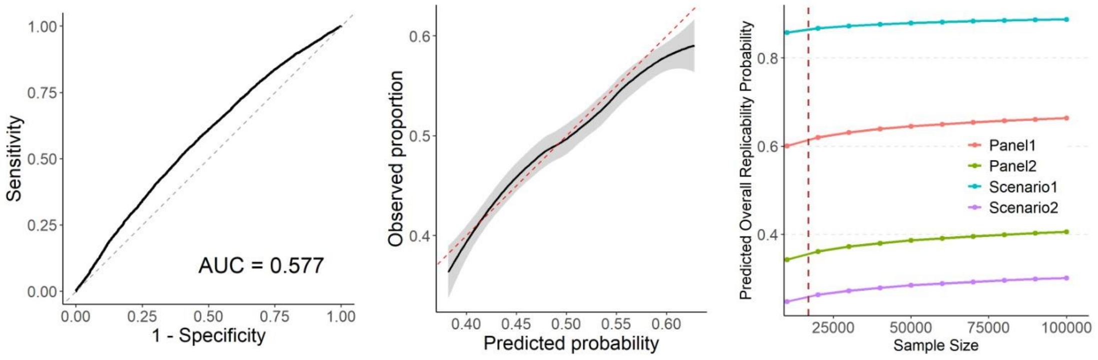


**Figure S10 | Predictive performance of logistic regression models for overall replicability.** AUC-ROC curves, calibration curves, and replicability prediction curves were shown for (**a**) - (**d**) two representative phenotypes (fluid intelligence and neuroticism) under standard and IPW logistic regression models.

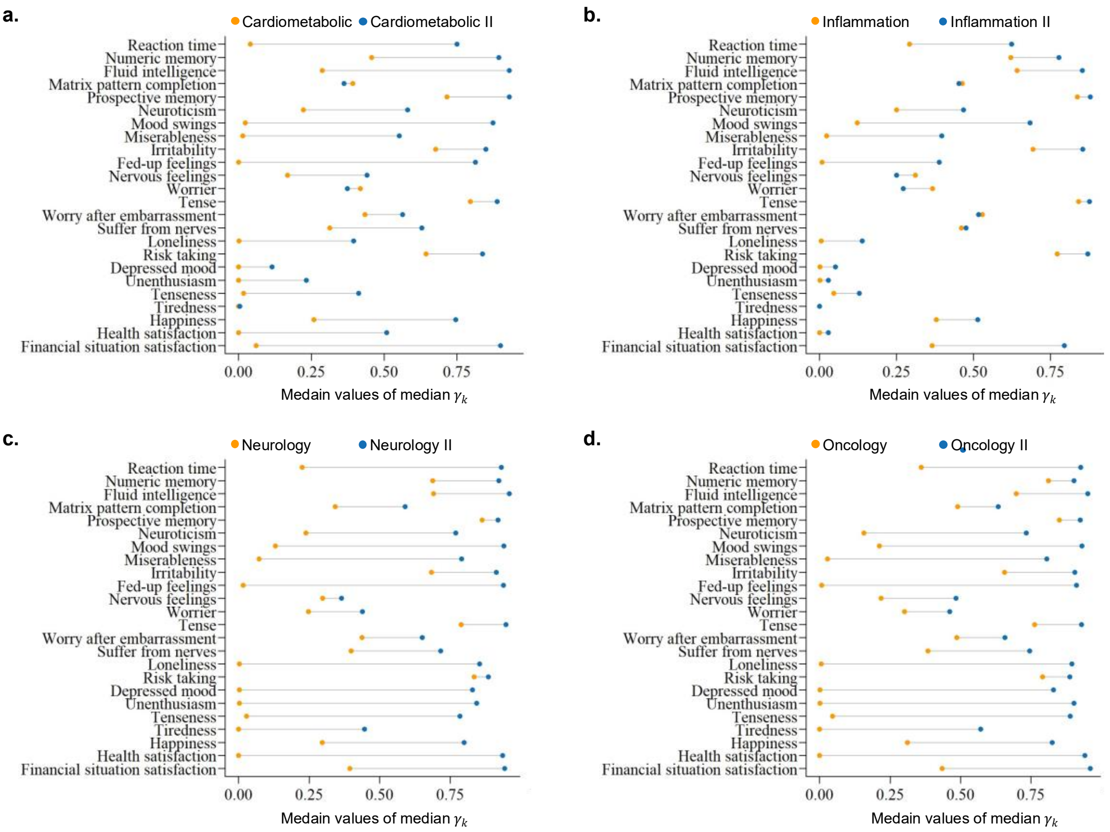


**Figure S11 | Individual replicability assessment for sub-panels.** Median values of the median individual irreplicability quantity $\gamma_k$ were compared between sub-panels from Panel 1 and Panel 2 for cognitive function and mental health traits. **a**, Cardiometabolic versus Cardiometabolic II. **b**, Inflammation versus Inflammation II. **c**, Neurology versus Neurology II. **d,** Oncology versus Oncology II. Each point represented the median value of median $\gamma_k$ for a given trait, with lower values indicating higher individual replicability.

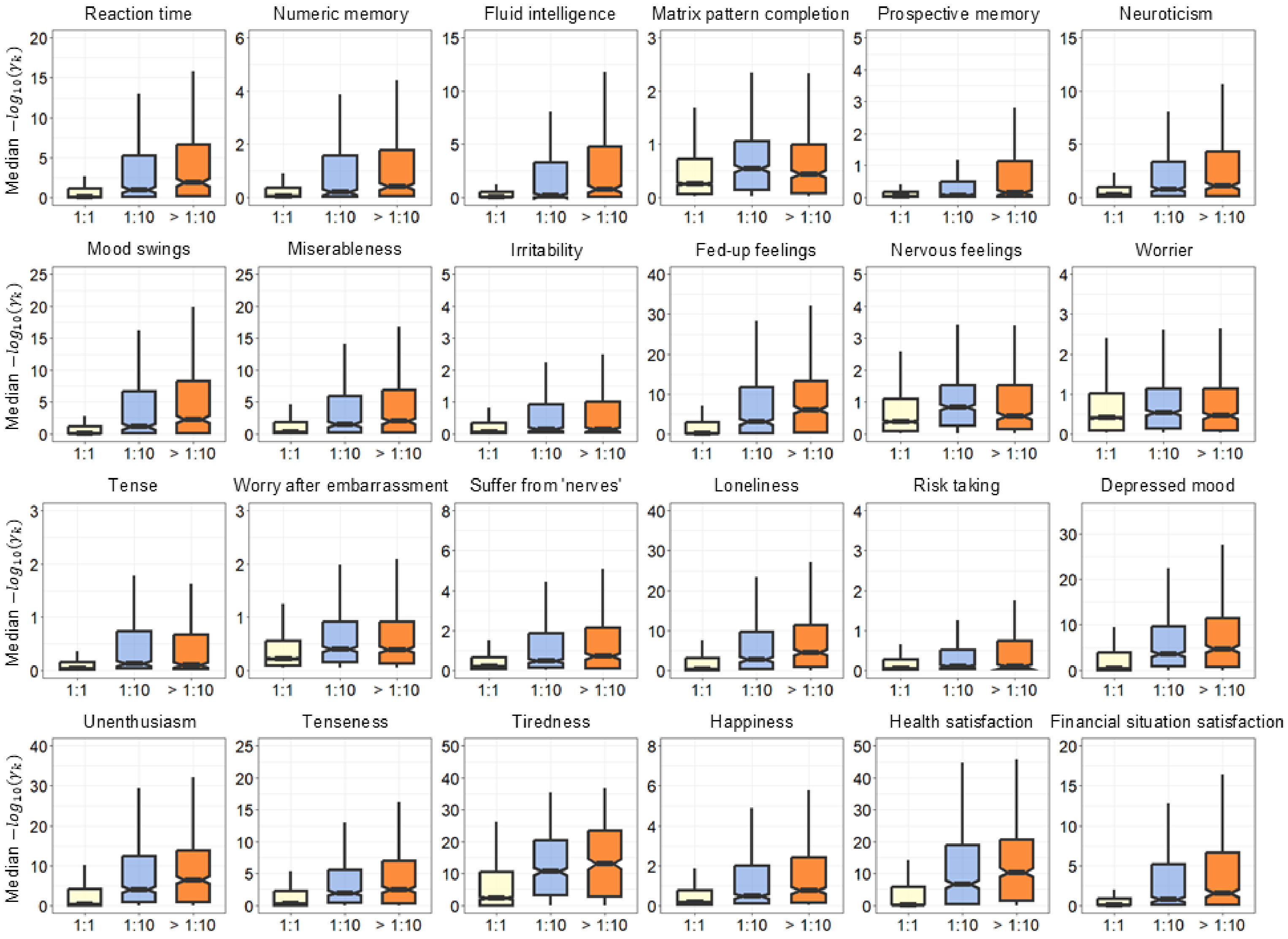
Reaction time
Numeric memory
Fluid intelligence
Matrix pattern completion
Prospective memory
Neuroticism
Mood swings
Miserableness
Irritability
Fed-up feelings
Nervous feelings
Worrier
Tense
Worry after embarrassment
Suffer from 'nerves'
Loneliness
Risk taking
Depressed mood
Unenthusiasm
Tenseness
Tiredness
Happiness
Health satisfaction
Financial situation satisfaction
Median $-log_{10}(\gamma_k)$
1:1
1:10
> 1:10

**Figure S12 | Individual replicability across dilution levels.** Boxplots of individual replicability (median $-\log_{10}(\gamma_k)$) by dilution levels. Each boxplot presented the median, lower- and upper-quartiles (Q1-Q3), with upper and lower whiskers representing 1.5x inter-quartile range. The x-axis represented different dilution levels and >1:10 represents more-abundant dilution levels (1:100 to 1:100,000).

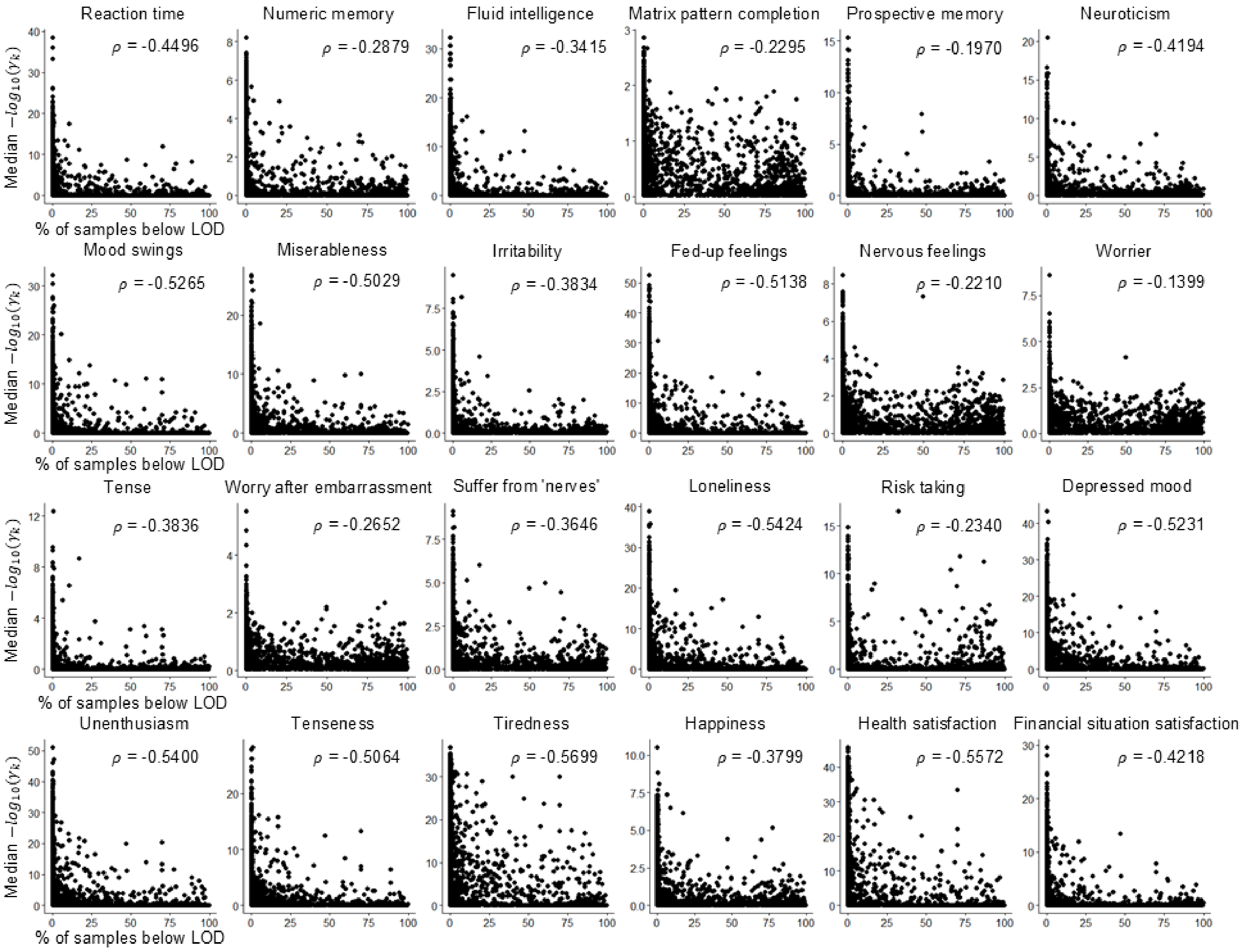
Reaction time
ρ = -0.4496
Numeric memory
ρ = -0.2879
Fluid intelligence
ρ = -0.3415
Matrix pattern completion
ρ = -0.2295
Prospective memory
ρ = -0.1970
Neuroticism
ρ = -0.4194
Mood swings
ρ = -0.5265
Miserableness
ρ = -0.5029
Irritability
ρ = -0.3834
Fed-up feelings
ρ = -0.5138
Nervous feelings
ρ = -0.2210
Worrier
ρ = -0.1399
Tense
ρ = -0.3836
Worry after embarrassment
ρ = -0.2652
Suffer from 'nerves'
ρ = -0.3646
Loneliness
ρ = -0.5424
Risk taking
ρ = -0.2340
Depressed mood
ρ = -0.5231
Unenthusiasm
ρ = -0.5400
Tenseness
ρ = -0.5064
Tiredness
ρ = -0.5699
Happiness
ρ = -0.3799
Health satisfaction
ρ = -0.5572
Financial situation satisfaction
ρ = -0.4218
Median $-log_{10}(\gamma_k)$
% of samples below LOD

**Figure S13 | Individual replicability across proportion of samples below the LOD.** Scatter plots of individual replicability (median $-\log_{10}(\gamma_k)$) by proportion of samples below the LOD. Each subfigure represented one trait, with the Spearman correlation coefficient ($\rho$) indicating the strength and direction of association.
